# Dissecting Bioelectrical Networks in Photosynthetic Membranes with Electrochemistry

**DOI:** 10.1101/2025.03.18.643929

**Authors:** Joshua M. Lawrence, Rachel M. Egan, Laura T. Wey, Karan Bali, Xiaolong Chen, Darius Kosmützky, Mairi Eyres, Lan Nan, Mary H. Wood, Marc M. Nowaczyk, Christopher J. Howe, Jenny Z. Zhang

## Abstract

Photosynthetic membranes contain complex networks of redox proteins and molecules which direct electrons along various energy-to-chemical interconversion reactions important for sustaining life on Earth. Analysing and disentangling the mechanisms, regulation and interdependencies of these electron transfer pathways is extremely difficult, owing to the large number of interacting components in the native membrane environment. Whilst electrochemistry is well established for studying electron transfer in purified proteins, it has proven difficult to directly wire into proteins within their native membrane environments, and even harder to probe on a systems-level the electron transfer networks they are entangled within. Here, we show how photosynthetic membranes from cyanobacteria can be directly wired to electrodes to access their complex electron transfer networks. Measurements of native membranes with structured electrodes revealed distinctive electrochemical signatures, enabling analysis from the scale of individual proteins to entire biochemical pathways, as well as their interplay. This includes measurements of overlapping photosynthetic and respiratory pathways, the redox activities of membrane-bound quinones, along with validation using *in operando* spectroscopic measurements. Importantly, we further demonstrated direct extraction of electrons from native membrane-bound Photosystem I at -600 mV, which is ∼1 V more negative than from purified photosystems. This finding opens up opportunities for biotechnologies for solar electricity, fuel and chemical generation. We foresee this electrochemical method being adapted to analyze other photosynthetic and non-photosynthetic membranes, as well as aiding the development of new biocatalytic, biohybrid and biomimetic systems.

## Introduction

The membrane-localized electron transfer chains (ETCs) of living organisms drive a diverse array of respiratory and photosynthetic processes, acting as the primary agents of energy flow in ecosystems^1,2^. Furthermore, ETCs and their components can be harnessed by researchers as electrocatalysts for the sustainable generation of electricity, fuels and high-value chemicals^3,4^. It is therefore essential to create tools for studying the molecular mechanisms underpinning the function of ETCs.

Analysis of ETCs is often complicated by the fact that they include various bifurcating electron transfer pathways comprised of many different redox proteins and cofactors^3,5,6^. Many organisms contain multiple ETCs within the same membrane, with these pathways often sharing components or catalysing coupled or opposing chemical reactions^7–11^. For example, photosynthetic membranes, such as the thylakoid membranes of cyanobacteria, contain highly complex networks of electron transfer reactions^7^, which limits our ability to analyze the activity, function and interactions of pathways and their components, which is needed to engineer improved productivity in those pathways. The main methodologies used to analyze electron transfer in biological systems are *in vitro* characterisation of purified redox proteins^*12–15*^, *in vivo* spectroscopy of highly-expressed redox proteins^*16–19*^, or indirect measurements on the impact of electron transfer processes on organism growth and physiology^20–23^. Methods that provide a systems-level understanding of ETC activity, whilst also being applicable to a wide variety of biomembranes, are lacking.

Electrochemistry has been widely applied to purified redox proteins and other biomolecules to analyze the mechanism and kinetics of their electron transfer reactions^13,14^, although biocomponents are typically in non-native environments. Similar electrochemical methods have been applied to biofilms of diverse microorganisms, which in the case of photosynthetic microorganisms have revealed complex electrochemical signatures^24–29^. Whilst these information-rich signatures have been shown to relate to the photosynthetic and respiratory activity of the cells, analysis has proven difficult owing to the many barriers separating the ETCs and the electrode^29^. Pioneering studies have investigated the interactions between biomembranes and electrodes, including many photosynthetic membranes^30–39^. However, these studies relied on genetic or chemical treatments which disrupt the native membrane environment. This has limited the ability of bioelectrochemistry to discriminate between different electron transfer pathways, hindering analytical applications. For this work, we hypothesized that with an optimized bio-electrode interface it should be possible to measure (previously missed) electrochemical signatures from native photosynthetic membranes directly, analysis of which could obtain information about their complex electron transfer pathways.

Herein, we established an analytical electrochemical approach to study the thylakoid membranes of cyanobacteria (Figure 1a). These photosynthetic membranes are the most protein dense in nature^40^ with perhaps the most complex network of interdependent electron transfer pathways (Figure 1b), including both a photosynthetic and respiratory ETC (PETC and RETC). Through the use of structured electrodes which provide an enhanced bio-electrode interface, sensitive photoelectrochemical signals were measured from even crudely isolated membranes. As hypothesized, distinctive electrochemical signatures could be resolved. Measurements of these signatures performed under various experimental conditions enabled analysis of biological electron transfer within these membranes from the scale of individual proteins to entire ETCs.

**Figure 1.**
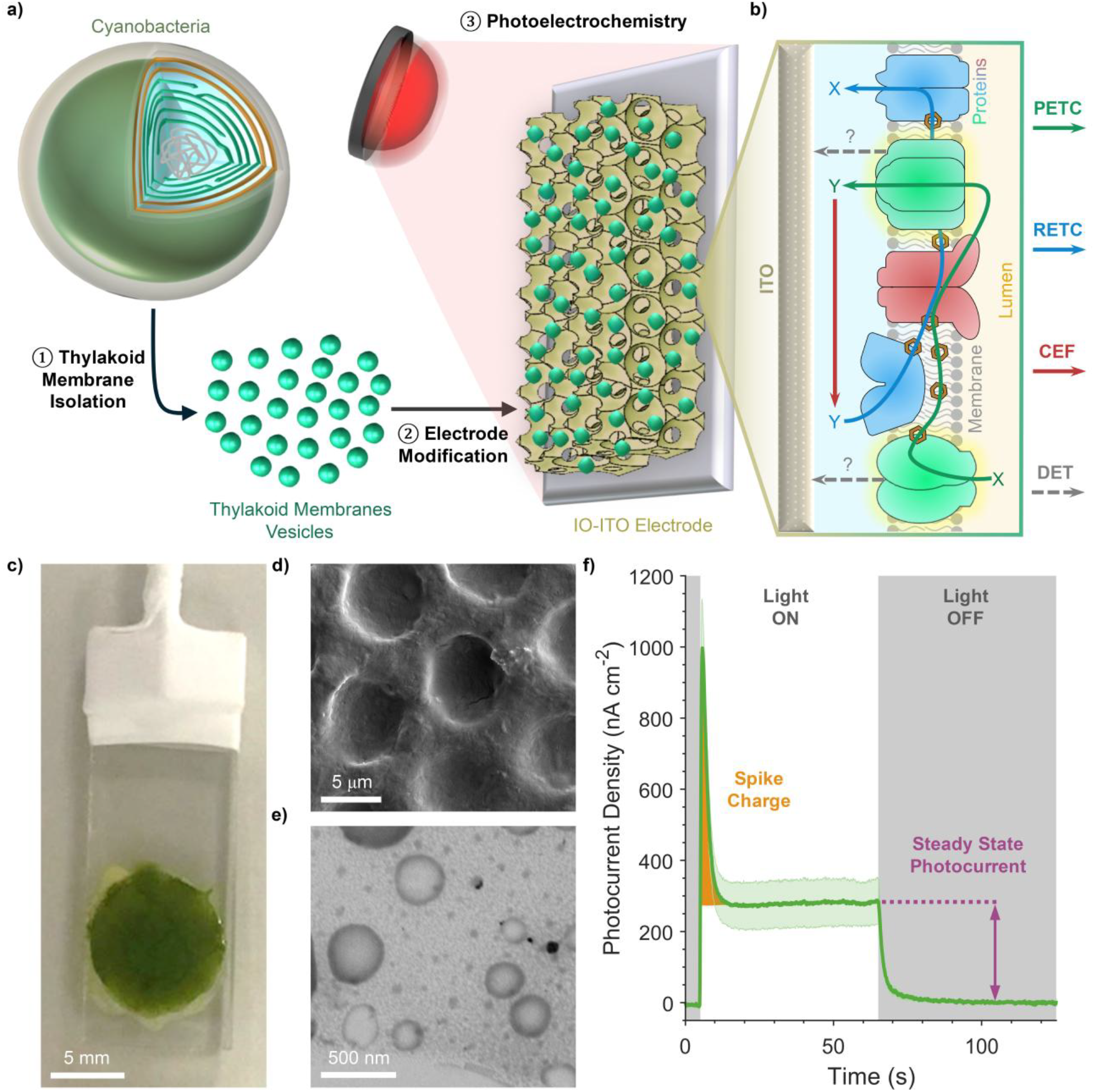
Biomembrane electrochemistry of cyanobacterial thylakoid membranes. **a)** Experimental workflow of thylakoid membrane electrochemistry experiments. **b)** Schematic depicting the electron transport network of cyanobacterial thylakoid membranes interfaced with the electrode. The thylakoid membrane electron transport pathways can be measured due to electron transfer between their components and the electrode. **c)** Picture, and **d)** SEM micrograph of a *Synechocystis* thylakoid membrane-modified electrode. **e)** STEM micrograph of isolated thylakoid membranes. **f)** Example thylakoid membrane photocurrent profiles, with key parameters labelled. All photocurrent measurements, including this, were performed under standard conditions (Methods), unless stated otherwise. Data presented as the mean of biological replicates ± S.E.M. (*n = 3*). CEF, cyclic electron flow; DET, direct electron transfer; IO-ITO, inverse opal-indium tin oxide; PETC, photosynthetic electron transport chain; RETC, respiratory electron transport chain.

## Results

### Wiring Thylakoid Membranes to Electrodes

For photosynthetic membranes samples, we used isolated thylakoid membranes from the model cyanobacterium *Synechocystis* sp. PCC 6803 (*Synechocystis* hcenterforth). This analyte was chosen due to its well-characterized molecular biology and availability of mutants, whilst still containing a highly complex network of biological electron transfer pathways. We developed a straightforward thylakoid membrane isolation method (Figure S1) which yielded membrane vesicles with a diameter of 100-250 nm and which maintained their *in vivo* topology and photosynthetic activity (Supporting Results 1, Figure S2-4). To ensure efficient wiring of membranes to the electrode, vesicles were adsorbed to hierarchically structured and translucent inverse opal-indium tin oxide (IO-ITO) electrodes^41^ with a pore size of 10 μm (Supporting Results 2, Figure S5-7).

Photocurrents of thylakoid membrane-modified electrodes were recorded in chronoamperometry experiments, where the change in current over a photoperiod was measured at a specific working electrode potential (*E*^app^). These measurements were performed under moderate red light (50 μmol photons m^-2^ s^-1^ at 680 nm) in a photoelectrochemical cell (Figure S8a-b). Data analysis was performed with bespoke software (Supporting Results 3, Figure S9, Materials and Methods). Notably, a distinct photocurrent profile was observed, consisting of a sharp spike in current in the dark-light transition, followed by relaxation to a steady state light current, with a rapid return to the steady state dark current following a light-dark transition (Figure 1f). These photocurrent profiles differ from those observed with individual photosystems (proteins which drive photosynthetic electron transport; which exhibit monophasic profiles), as well as those of cyanobacterial biofilms (which have more complex profiles)^29,41^. The features were quantified as two parameters: (i) the ‘Spike Charge’ (the charge contained within the initial spike feature); and (ii) the ‘Steady State Photocurrent’ (the positive difference in current between the light and dark steady states). This enabled quantitative analysis of these parameters under different experimental conditions.

Experimental conditions were optimized to enable thylakoid membrane photocurrents to be recorded in physiologically relevant conditions (Figure S10), which led to the selection of standard conditions which were used in subsequent electrochemistry experiments (Supporting Results 4, Materials and Methods).

### Probing Photosynthetic Electron Transport Pathways

Cyanobacteria thylakoid membranes contain a complete PETC (Figure 2a), which begins with photoexcitation by photosystem II (PSII) using electrons obtained from water oxidation, with these electrons subsequently being transported via the plastoquinone/plastoquinol (PQ/PQH_2_) pool to cytochrome *b*_6_*f*, plastocyanin, and photosystem I (PSI). PSI performs an additional photoexcitation, followed by electron transfer to ferredoxin and FNR, which synthesises the redox carrier NADPH^7^. The different dependencies of the Steady State Photocurrent and Spike Charge on chl *a* loading, light intensity and pH (Supporting Results 4, Figure S10a-c,) led us to hypothesize that these two photocurrent parameters originate from different direct electron transfer (DET) pathways between the thylakoid membranes and the electrode (Figure 1b), which may provide information on electron transfer through components of the PETC. To test this, photocurrents were recorded in the presence of different inhibitors and substrates, light conditions and working electrode potentials (*E*_app_) to discriminate between electron transfer pathways from the PETC to the electrode.

**Figure 2.**
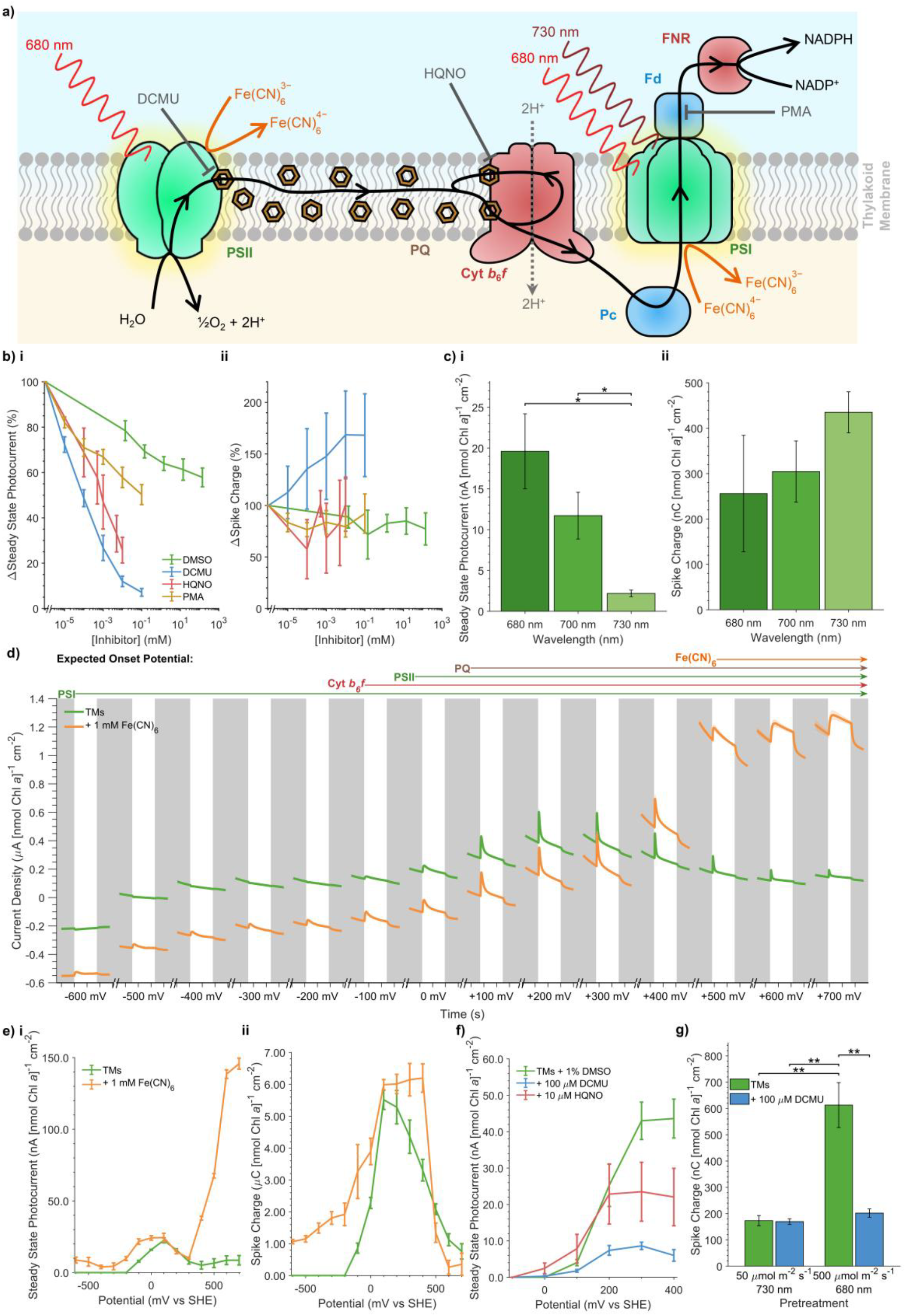
Electrochemical measurements of the photosynthetic electron transport chain. **a)** Schematic depicting the photosynthetic electron transport chain of *Synechocystis* thylakoid membranes. The sites of action of inhibitors, electron mediators and wavelengths of light are shown. **b)** Effect of various inhibitors of photosynthetic electron transport on the relative change of photocurrent parameters. DMSO alone is included as a control. **c)** Effect of illumination wavelength on photocurrent parameters. **d)** Stepped chronoam-perometry scan of thylakoid membranes, with each successive photocurrent recorded at an increasing *E*_app_ value (mV vs SHE) and in the absence and presence of 1 mM Fe(CN)_6_ along with an enzymatic oxygen removal systems (1 mM glucose, 100 μg mL^-1^ glucose oxidase, 50 μg mL^-1^ catalase). X-axis tick spacings represent 20 s of experimental time. **e)** Photocurrent parameters calculated from **d. f)** Effect of *E*_app_ and photosynthetic inhibitors on Steady State Photocurrents, calculated from stepped chronoamperometry experiments performed without enzymatic oxygen removal. **g)** Effect of light pre-treatment conditions on Spike Charge, calculated from photocurrents recorded after 10 s of pretreatment followed by 10 s of darkness. Supporting data in Figure S10. Sub-panels depict measurements of **(i)** Steady State Photocurrent and **(ii)** Spike Charge magnitudes. All data presented as the mean of biological replicates ± S.E.M. (n = 3). p-values calculated using two-tailed unpaired t-tests (*** p≤0.001, ** 0.001<p≤0.01, p≤0.05). Chl, chlorophyll; Cyt b6f, cytochrome b6f; DCMU, 3-(3,4-dichlorophenyl)-1,1-dimethylurea; DMSO, dimethyl sulfoxide; Fd, ferredoxin; 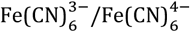, ferri-/ferrocyanide redox couple; FNR, ferredoxin-NADP+ reductase; HQNO, N-oxo-2-heptyl-4-Hydroxyquino-line; Pc, plastocyanin; PQ, plastoquinone; PSII, photosystem II; PSI, photosystem I; NADP(H), nicotinamide adenine dinucleotide phosphate; PMA, phenylmercuric acetate; SHE, standard hydrogen electrode; TM, thylakoid membrane.

Titrations of several PETC inhibitors (Figure 2a) were performed to determine their inhibition of photocurrent parameters relative to a DMSO control (Figure 2b). 3-(3,4-Dichlorophenyl)-1,1-dimethylurea (DCMU), a competitive inhibitor of the PSII Q_B_ site^42^, provided a strong inhibition of Steady State Photocurrent up to 93% but had no inhibitory effect on the Spike Charge, suggesting that whilst Steady State Photocurrent is PSII-dependent, Spike Charge is purely PSI-dependent. DCMU also provided a small enhancement in the Spike Charge, although this could be caused by the signal from the Steady State Photocurrent blocking the spike feature in the absence of DCMU. 2-Hep-tyl-4-hydroxyquinoline N-oxide (HQNO) is a competitive inhibitor of the cytochrome *b*_6_*f* Q_n_ site^43^ which prevents electron transfer downstream of haem *b*_n_ without effecting electron transfer to PSI via plastocyanin^44^. HQNO did not inhibit the Spike Charge, consistent with this feature being PSI-dependent, whilst inhibiting the Steady State Photocurrent up to 74%, suggesting a sizeable proportion of this feature originates from DET between cytochrome *b*_6_*f* and the electrode. Phenylmercuric acetate (PMA), a ferredoxin inhibitor^45^, had no effect compared to the DMSO control. This is consistent with ferredoxin being lost during thylakoid membrane isolation (Supporting Results 1, Figure S4f) and PSI performing electron transfer to the electrode. Photocurrents recorded with illumination at longer wavelengths (700 and 730 nm) which selectively excite PSI over PSII^46^ further showed the PSI-dependency of the Spike Charge and PSII-dependency of the Steady State Photocurrent (Figure 2c).

In bioelectrochemistry experiments, the *E*_app_ of the working electrode can be used to control which cofactors can transfer electrons to the electrode based on their midpoint potential (*E*_m_). We exploited this in stepped chronoamperometry experiments, where an enzymatic oxygen removal regime was used to enable recordings of photocurrents at increasing *E*_app_ values from -600 to +700 mV vs standard hydrogen electrode (SHE)^47^. The most negative *E*^app^ at which photocurrent features appeared was used to distinguish between electron transfer from different PETC redox cofactors to the electrode (Figure 2d-e). The potential for the appearance of anodic (positive) photocurrents was observed at -100 mV vs SHE, with the Steady State Photo-current reaching its maximum at +100 mV vs SHE, consistent with the feature being dependent on DET from PSII and cytochrome *b*_6_*f* to the electrode^3,41^. A fast decline in the Spike Charge was observed at *E*_app_ > +200 mV vs SHE, which can be explained by the oxidation of PQH_2_ by the electrode^48^ (see below) competing with reduction of PSI. The ferri-/ferrocyanide redox couple has an *E*_m_ of +420 mV vs SHE (Figure S11); at *E*_app_ values below this it acts as a PSI electron donor^49^, whilst above this it acts as a PSII and cytochrome *b*_6_*f* electron acceptor^50,51^. Addition of this molecule led to the appearance of anodic photocurrents with a clear spike feature at *E*_app_ values as low as -600 mV vs SHE (Figure S12), the appearance of currents at such potentials can only be explained by DET from either PhQ or Fe_X_ of PSI to the electrode^52^. Even at more positive *E*_app_ values where other electron transfer pathways to the electrode could predominate, we find no evidence for mediated electron transfer to the electrode in the absence of an exogenous mediator (Supporting Results 5, Figure S13-17). At *E*_app_ values ≥ +500 mV vs SHE a clear enhancement in the Steady State Photocurrent was observed in the presence of ferricyanide (Figure 2d-e), caused by mediation from PSII to the electrode. In this condition the Spike Charge disappeared entirely-consistent with the Steady State Photocurrent being caused by PSII- and cytochrome *b*_6_*f*-dependent electron transfer and the Spike Charge by PSI-dependent electron transfer.

To identify the redox cofactors responsible for electron transfer to the electrode, stepped chronoamperometry was performed in the presence of inhibitors. Steady State Photocurrent inhibition was only observed at *E*_app_ values ≥ +100 mV and +300 mV vs SHE for DCMU and HQNO respectively (Figure 2f), consistent with electron transfer from PSII Q_B_ and cytochrome *b*_6_*f* haem *c*_n_ to the electrode. Cyclic voltammetry, an electrochemical technique which can be used to measure the *E*_m_ of cofactors^13^, revealed the presence of a cofactor with an *E*_m_ of +77 mV vs SHE consistent with PQ(H_2_) (Supporting Results 6, Figure S19). This suggests that at more positive *E*_app_ values PQH_2_ could be oxidized by the electrode, potentially disrupting electron transfer from PSII to PSI (Figure 2a). To test if this occurs under standard conditions, photocurrents were recorded following a pre-treatment with either 500 μmol photons m^-2^ s^-1^ of 680 nm light (to promote PQ reduction by PSII), or 50 μmol photons m^-2^ s^-1^ of 730 nm light (to promote PQH_2_ oxidation by PSI) (Figure S19). A 3.5-fold enhancement in the Spike Charge was observed in the former condition, with DCMU abolishing this enhancement (Figure 2g), demonstrating how PSII activity enhances PSI-dependent electron transfer to the electrode and confirming functional electron transfer through the PETC. Taken together, these results demonstrate that biomembrane electrochemistry enables measurement of electron transfer through multiple components of the cyanobacterial PETC (Figure 4).

### Probing Respiratory Electron Transport Pathways

Cyanobacterial thylakoid membranes, unlike those found in plant and algal chloroplasts, contain a complete RETC_7_ (Figure 3a). This RETC contains various dehydrogenases which oxidize electron donors (such as ferredoxin, NAD(P)H and succinate) and reduce PQ, as well as terminal oxidases which oxidize PQH_2_ (directly or via plastocyanin) and use oxygen as a terminal electron acceptor. The cyanobacterial PETC and RETC share components with each other, giving rise to cyclic electron flow which connects the pathways in a complex electron transfer network_7_ (Figure 2a). This makes it especially hard to disentangle the activity of the RETC from that of the PETC in cyanobacterial thylakoid membranes.

**Figure 3.**
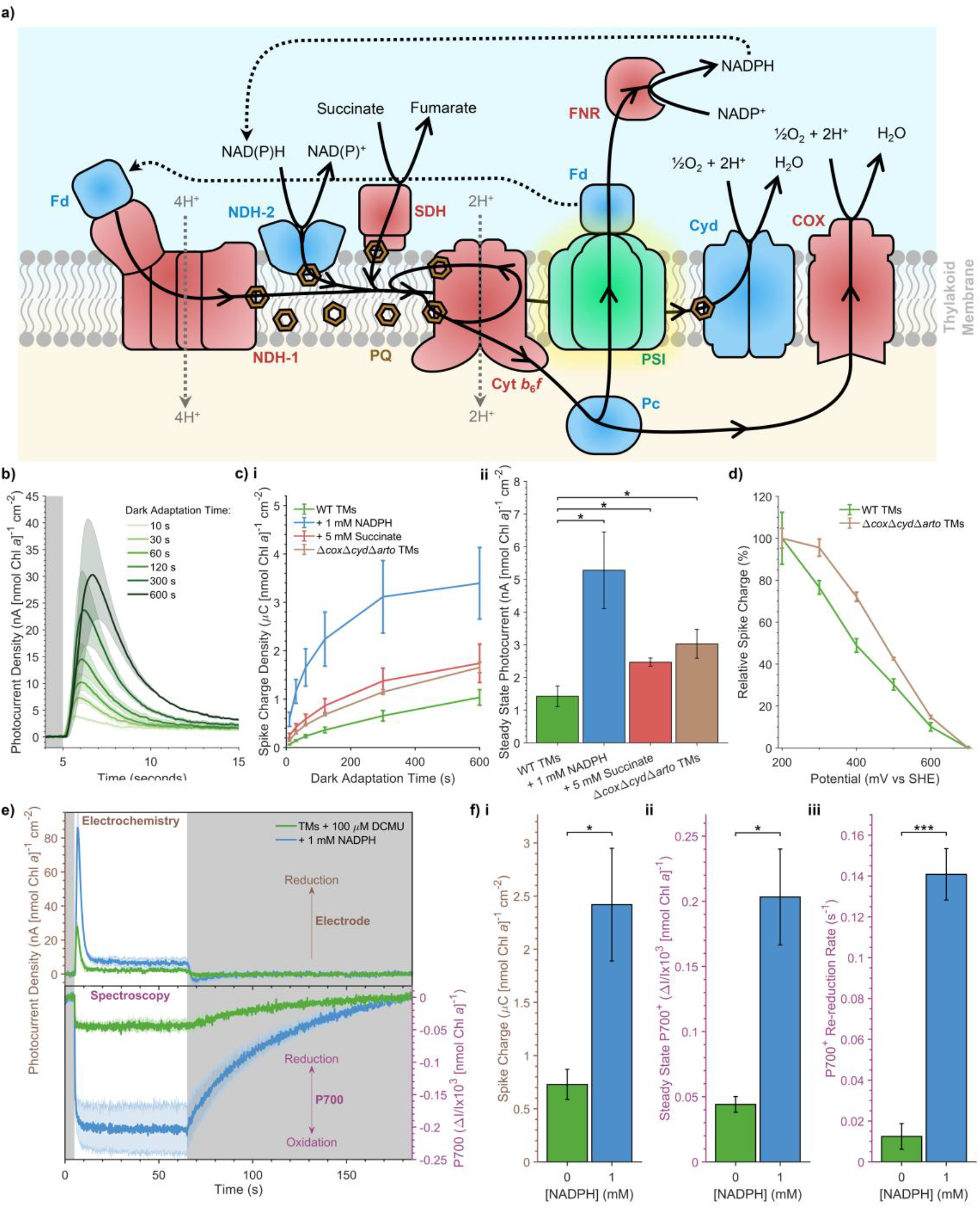
Electrochemical measurements of the respiratory electron transport chain. **a)** Schematic depicting the respiratory electron transport chain of *Synechocystis* thylakoid membranes. Cyclic electron flow around PSI is also depicted. **b)** Thylakoid membrane photocurrent profiles recorded in increasing dark adaptation times. **c)** The effect of dark adaptation time on photocurrent parameters recorded in conditions with a more reduced plastoquinone pool. This was achieved by addition of substrates which reduce plastoquinone via dehydrogenases (succinate and NADPH) or by utilising a mutant lacking respiratory terminal oxidases which oxidize the plastoquinone pool (Δ*cyd*Δ*cox*Δ*arto*). Sub-panels depict measurements of **(i)** Spike Charge (at all dark adaptation times) and **(ii)** Steady State Photocurrents (at a dark adaptation time of 60 s). **d)** The relative decay in Spike Charge measured at *E*_app_ values ≥ +200 mV vs SHE for thylakoid membrane from wild type and Δ*cyd*Δ*cox*Δ*arto* cells without enzymatic oxygen removal. **e)** *In oper- ando* measurements of thylakoid membrane photocurrents alongside the oxidation of the P700 reaction center of photosystem I, measured in the absence and presence of 1 mM NADPH. **g)** Measurements of **(i)** Spike Charge, **(ii)** steady state P700 oxidation, and **(iii)** P700 re-reduction rate calculated from data in panel **f**. All experiments performed in the presence of 100 μM DCMU. All data presented as the mean of biological replicates ± S.E.M. (*n = 3*). *p*-values calculated using two-tailed unpaired *t*-tests (*** *p*≤0.001, ** 0.001<*p*≤0.01, *p*≤0.05). Chl, chlorophyll; COX, cytochrome *c* oxidase; Cyd, cytochrome *bd* oxidase; Cyt *b*_6_*f*, cytochrome *b*_6_*f*; Fd, ferre-doxin; DCMU, 3-(3,4-dichlorophenyl)-1,1-dimethylurea; FNR, ferredoxin-NADP_+_ reductase; NAD(P)(H), nicotinamide adenine dinucleotide (phosphate); NDH-1, type I NAD(P)H dehydrogenase; NDH-2, type II NAD(P)H dehydrogenase; Pc, plastocyanin; PQ, plas-toquinone; PSI, photosystem I; SDH, succinate dehydrogenase; SHE, standard hydrogen electrode; TM, thylakoid membrane; WT, wild type.

**Figure 4.**
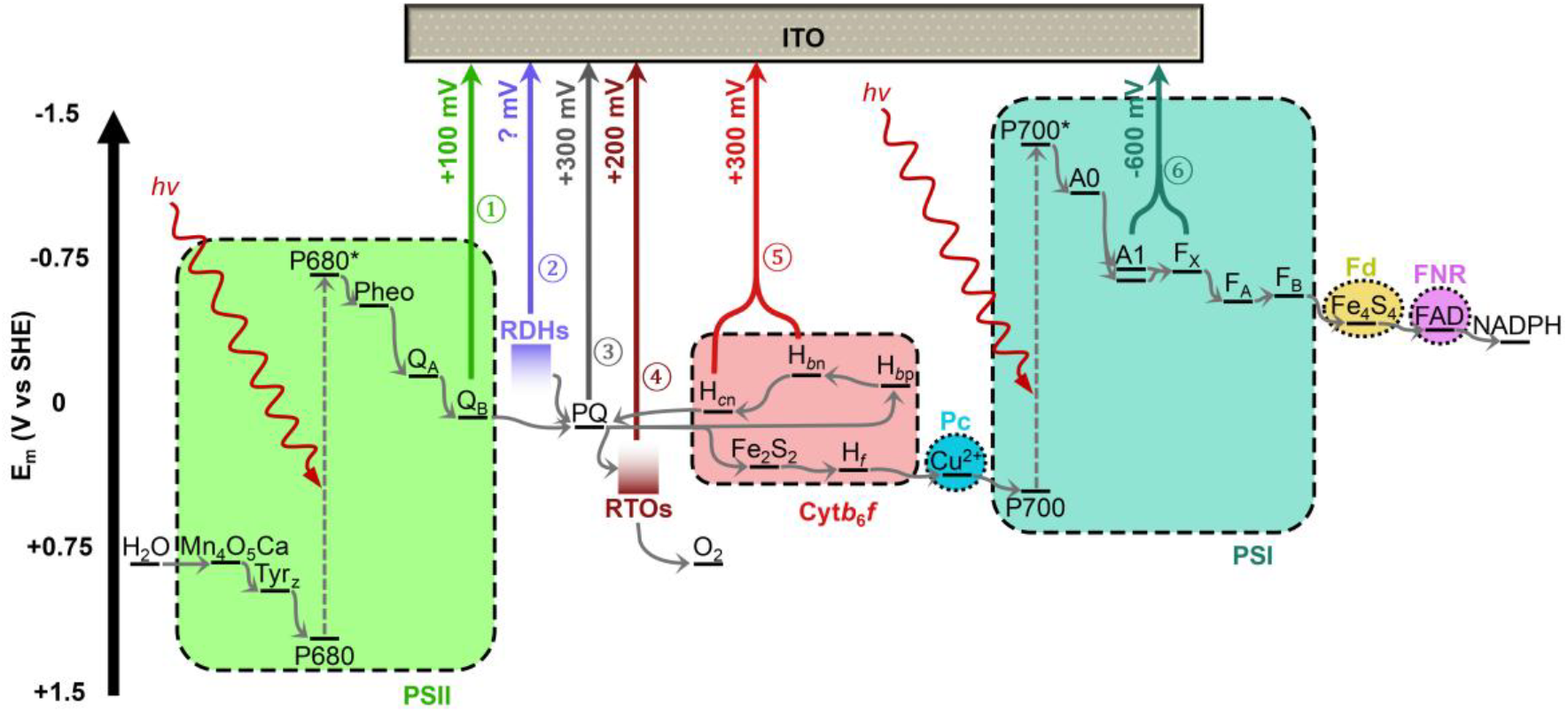
Model of interfacial electron transfer pathways. Identified thylakoid membrane-electrode electron transport pathways. Redox potentials^3^ and suspected routes of electron transport between redox cofactors and the electrode are shown alongside the lowest *E*_app_ values these pathways were observed at. Supporting information in Supplementary Table 1. A, acceptor; Cyt *b*_6_*f*, cyto-chrome *b*_6_*f*; F, Fe-S cluster; FAD, flavin adenine dinucleotide; Fd, ferredoxin; FNR, ferredoxin-NADP^+^ reductase; H, haem; ITO, indium tin oxide; NADP(H), nicotinamide adenine dinucleotide phosphate; P, primary donor; Pc, plastocyanin; Pheo, pheophytin; PQ, plastoquinone; PSII, photosystem II; PSI, photosystem I; Q, quinone; RDHs, respiratory dehydrogenases; RTOs, respiratory terminal ox-idases; SHE, standard hydrogen electrode; Tyr, tyrosine.

Our findings that the Spike Charge feature was linked to both the NADPH concentration (Figure S18) and the PQ pool redox state (Figure 2g) led us to hypothesize that it was dependent on RETC activity. In this model, reduction of PSI in the dark by the RETC (Figure 3a) leads to the accumulation of charge which upon illumination is rapidly dissipated via electron transfer from PSI to the electrode, thereby giving rise to the spike in the current. To test this hypothesis, photocurrents were recorded in the presence of DCMU and using 700 nm light to excite PSI selectively, with varying dark adaptation times (the length of the dark period prior to illumination) used to control the extent of PQ reduction by the RETC. A clear increase in the Spike Charge was observed at increasing dark adaptation times (Figure 3b), suggesting the Spike Charge encodes information on the RETC activity occurring in the preceding dark period. The same experiments were performed in the presence of electron donors for PQ-reducing dehydrogenases (Figure 3a). Addition of NADPH and succinate both provided enhancements in the Spike Charge at all dark adaptation times (Figure 3ci and S20a-b). This activity can be attributed to the action of NAD(P)H dehydrogenases in the case of NADPH and succinate dehydrogenase in the case of succinate_7_. Furthermore, enhancements in the Spike Charge were also observed in thylakoids obtained from mutants lacking all PQH_2_-oxidising terminal oxidases (Δ*cox*Δ*cyd*Δ*arto*), which have previously been shown to have a more reduced PQ pool^23,53^. Steady State Photocurrent enhancements independent of the dark adaptation time were also observed in these conditions (Figure 3cii and S20c), demonstrating an enhancement of steady state PSI activity in conditions with a more reduced PQ pool. These results provide a clear demonstration that biomembrane electrochemistry can be used to measure RETC in cyanobacterial thylakoid membranes. Furthermore, photocurrents obtained with thylakoid membranes from species other than *Synechocystis* suggest these PQ-reducing pathways may be specific to cyanobacteria^2,7^ (Supporting Results 7, Figure S21).

Addition of NADPH and succinate to the electrolyte were also found to provide an enhancement of the dark current (Figure S20d), which could be caused by dehydrogenase enzymes being wired to the electrodeeither directly or via PQ(H_2_). Whilst this could be influenced by abiotic factors (such as changes in capacitance^13^), electron transfer from RETC components to the electrode is feasible. For example, some of the decay in the Spike Charge magnitude observed at *E*_app_ values > +200 mV vs SHE (Figure 2d-e and S14) could be caused by the electrode oxidising PQH_2_ via terminal oxidases (Figure 3a) rather than directly. To test this *E*_app_-dependent changes in the Spike Charge were measured using thylakoid membranes from WT and Δ*cox*Δ*cyd*Δ*arto* cells in the presence of DCMU. Whilst both conditions exhibited a decay in Spike Charge at more positive *E*_app_ values, this decay was slower in the Δ*cox*Δ*cyd*Δ*arto* mutant, consistent with diminished oxidation of PQH_2_ by the electrode (Figure 3d and S22). Furthermore, the start of the decay was shifted from +200 to +300 mV vs SHE in the mutants. These results can be explained by electron transfer from respiratory terminal oxidases to the electrode at *E*_app_ ≥ +200 mV vs SHE, with direct oxidation of PQH_2_ occurring at *E*_app_ ≥ +300 mV vs SHE-a large overpotential consistent with previous electrochemical measurements performed with synthetic membranes^48^. These results demonstrate measurements of individual ETC components can be performed with biomembrane electrochemistry (Figure 4).

Spectroelectrochemistry measurements were performed to ensure that these electrochemical measurements match those obtained with established spectroscopic methods. A spectroelectrochemical cell was constructed (Figure S8c) for use in a Joliot-type spectrophotometer^17^, enabling in operando measurements of photocurrents and changes in the oxidation state of the P700 reaction center of PSI^54^ (Figure 3e-f). The peak of the Spike Charge feature in photocurrent measurements was correlated with the peak of P700 oxidation, further proving the Spike Charge is dependent on PSI electron transfer to the electrode. Addition of NADPH led to an increase in the oxidation of P700, consistent with it being more reduced prior to illumination due to reduction of the PQ pool by the RETC. The fold change of the Spike Charge in the presence and absence of NADPH was similar to that of the steady state P700 oxidation (3.3- and 4.6-fold respectively) suggesting the two parameters are equivalent to one another. Furthermore, the rate of P700 re-reduction in the dark was increased in the presence of NADPH by 11-fold, also consistent with a faster reduction of the PQ pool by the RETC. These results demonstrate that our electrochemical analysis of biomembranes gives results consistent with established biophysical techniques for analysing thylakoid membrane electron transfer, complementing these methods by enabling measurement of electron transfer at multiple points of the ETC (Figure 4).

## Discussion

Our results enable us to construct a model detailing the electron transfer pathways occurring across the bio-electrode interface (Figure 4, Table S1), with DET being the most likely mechanism of interfacial electron transfer (Supporting Results 5-6). These results are aligned with our understanding of cyanobacterial thylakoid membrane electron transfer *in vivo*_7_ (Supporting Discussion), establishing our approach of performing electrochemistry on membranes as an effective analytical method, even whilst using simply prepared thylakoid membrane samples. New observations, including the low *E*_app_ required for anodic PSI photocurrents (∼1V more negative than previously observed^55,56^) and the electrochemical measurements of PQ pool reduction in native membranes are, to our knowledge, new milestones.

Here we show that with optimized bio-electrode interfaces and controlled experimental conditions, it is possible to analyze electron transfer events using simple and reproducible electrochemical experiments. This is achieved without the need for protein overexpression or lengthy purification protocols, and the technique can be readily coupled to spectroscopic methods (Figure 3e-h). As such, this is a powerful yet highly accessible approach for analysing membrane electron transfer networks (Figure S23).

Furthermore, given the complexity of the cyanobacterial thylakoid membrane electron transfer network^7^ (Figure 1b) we expect that this biomembrane electrochemistry technique could be readily applied to other photosynthetic (Supporting Results 7) or non-photosynthetic membranes (such as those derived from mitochondria). We foresee immediate applications in characterising and engineering ETCs for fundamental research, enhancing photosynthetic yields, identifying the mechanism of inhibitor/herbicide interactions^57,58^, and informing biocatalytic and biomimetic redox cascade systems for sustainable chemical production^3,59,60^.

## ASSOCIATED CONTENT

### Supporting Information

Supporting Information is available free of charge at https://pubs.acs.org. It includes detailed materials and methods, supporting results (biological sample characterisation, electrode characterisation, protein structure analysis, additional electrochemical data) and supporting discussion.

### Supporting Data and Code

All raw data and analysis code is available free of charge at https://github.com/JLaw-rence96/Dissecting-bioelectrical-networks-in-photosyn-thetic-membranes-with-electrochemistry.

## AUTHOR INFORMATION

## Author Contributions

All authors have given approval to the final version of the manuscript.

## Notes

The authors declare no competing financial interest.

## ACKNOWLEDGMENT

The authors are grateful for support from the Biotechnology and Biological Sciences Research Council (BB/M011194/1 to J.M.L., BB/R011923/1 to J.Z.Z. & X.C.); the Worshipful Company of Leathersellers (J.M.L.); Trinity Hall, Cambridge (J.M.L.); the Novo Nordisk Foundation (NNF22OCOO79717 to L.T.W.); the Cambridge Trust (L.T.W., R.M.E. & L.N.), the Engineering and Physical Sciences Research Council (EP/R513180/1 to K.B.); Algae-UK (M.E.); the Chinese Scholarship Council (no. 202208060220 to L.N.); the Gates Cambridge Trust (D.G.K.); the Benn W Levy Trust (D.G.K.); and the Human Frontiers Science Program (LT000307/2019 to M.H.W.). The authors would like to thank Nicholas Plumeré and Henry Lloyd-Laney for helpful discussions.

