## Supporting Information for "Dissecting Bioelectrical Networks in Photosynthetic Membranes with Electrochemistry"

### Materials and Methods

#### Strains and growth conditions

Cyanobacterial strains (*Synechocystis* sp. PCC 6803 and *Synechococcus elongatus* PCC 7942) were grown photoautotrophically in BG-11 media supplemented with 10 mM bicarbonate[1]. The microalga *Chlamydomonas reinhardtii* cw15 (cc-1883) was grown photomixotrophically in TAP medium[2]. Solid cultures used 1.5 % agar (w/v) in the culture medium. Liquid cultures were grown from single colonies cultures under 40 µmol photons $m^{-2}$ $s^{-1}$ at 30 °C, with shaking of 120 rpm and growth tracked by the attenuance at 750 nm ($OD_{750}$).
The plant *Nicotiana benthamiana* was grown under a 16 hr white light (100 µmol photons $m^{-2}$ $s^{-1}$ ) / 8 hr dark photoperiod at 21 °C with regular watering. Seeds were stored at 4 °C prior to plantation. Plants were initially germinated from seeds for two weeks, after which seedlings were transferred to individual pots and grown for a further four weeks.

#### Thylakoid membrane isolation

Cyanobacterial membranes enriched in thylakoids were isolated using a previously described method with numerous modifications[3]. One batch provided sufficient sample for ~3 electrodes. Cyanobacterial cultures were grown to mid-stationary phase ($OD_{750}$ = 1.5±0.2), with 50 mL pelleted by centrifugation (3000 x g, 10 min) after which the supernatant was removed and discarded. Cell pellets were resuspended in 5 mL of lysis buffer (50 mM 2-(N-Morpholino)ethanesulphonic acid-NaOH (MES-NaOH), 5mM EDTA, 2mM aminocaproic acid, 1mM phenylmethylsulphonyl fluoride, 1X concentration of Roche cOmplete protease inhibitor cocktail, adjusted to pH 6.5) at room temperature and re-centrifuged (3000 x g, 10 min). All subsequent steps were performed at 4 °C away from direct light. The resulting supernatant was removed and discarded, whilst the pellet was resuspended in 900 µL of chilled lysis buffer and transferred to a 2 mL safe-lock microcentrifuge tube. Acid washed glass beads (425-600 µm) were added up to a total volume of 1.75 mL. Cell lysis was performed by two rounds of vortexing on maximum speed for 90 s, followed by a 60 s incubation on ice. The resulting solution was transferred to a 2 mL microcentrifuge tube. The remaining glass beads were washed thrice with 400 µL of lysis buffer (using a 1000 µL pipette tip with the end cut off), with each wash being transferred to the new microcentrifuge tube which was centrifuged to pellet unbroken cells (9000 x g, 1.5 min). The supernatant (lysate) was transferred to a 2 mL microcentrifuge tube and centrifuged (15,000 x g, 15 min). The supernatant (soluble fraction) was transferred to a brown 15 mL centrifuge tube, with 10 µL being removed and diluted into 990 µL of lysis buffer to determine the C-phycocyanin content. 3 kDa-filtered soluble fraction was prepared by centrifuging through a 3 kDa MWCO Amicon® Ultra-15 Centrifugal Filter Unit (5,000 x g, 1 hour). The remaining pellet (thylakoid membranes) was washed without resuspending in 200 µL of lysis buffer and centrifuged (15,000 x g, 15 min). The supernatant was then discarded and the pellet (thylakoid membranes) resuspended in 20 µL of storage buffer (50 mM Tris-HCl, 1M sucrose, adjusted to pH 6.5) using a fine paintbrush and transferred to a dark. This thylakoid membrane suspension was transferred to a brown 500 µL microcentrifuge tube, with 1 µL being removed and diluted into 999 µL of methanol to determine the chl *a* concentration. The thylakoid membrane suspension was then diluted with storage buffer to a final chl *a* concentration of 2 µg $\mu L^{-1}$, unless otherwise stated.

Green algal thylakoid membranes were isolated from 150 mL of *C. reinhardtii* culture grown to mid-exponential phase ($OD_{750}$=0.9±0.05) using a previously developed extraction method[4]. One batch prepared according to this method provided sufficient thylakoid membranes for ~3 electrodes. Following extraction, 1 µL of thylakoid membrane suspension was diluted in 999 µL of a 4:1 acetone to methanol mixture to determine the chl *a*+*b* concentration. The thylakoid membrane suspension was then diluted to a final chl *a*+*b* concentration of 750 ng $\mu L^{-1}$.

Plant thylakoid membranes were isolated from 4 g of *N. benthamiana* leaves using extraction method A2 from Chen *et al*[5]. One batch prepared according to this method provided sufficient thylakoid membranes for ~20 electrodes. Following extraction, 1 µL of thylakoid membrane suspension was diluted in 999 µL of a 4:1 acetone to methanol mixture to determine the chl *a*+*b* concentration. The thylakoid membrane suspension was then diluted to a final chl *a*+*b* concentration of 4 µg $\mu L^{-1}$.

#### Electrode preparation

IO-ITO electrodes for electrochemistry experiments were made according to a previously developed protocol with modifications[6]. Fluorine-doped tin oxide-coated glass (Merck, 735167, 7 Ω $sq^{-1}$) cut into 1 cm x 3 cm rectangles was used as a support. These were first cleaned by bath sonication in isopropanol and then in ethanol. After drying in an oven at 100 °C for 1 hr, circular stencils of 1 cm diameter were made on the conductive face with electrical tape. The slides were UV-ozone treated using a Bioforce Nanosciences UV-Ozone Procleaner for 15 min. 1 mL of a 2.5% (w/w) solution of polystyrene beads (AlfaAesar, 42717, 10 µm) was centrifuged (15,000 x g, 4 min), with 750 µL of the supernatant being discarded. The pellet was then resuspended in the remaining supernatant, with 30 µL of the resulting solution being dropcast into the circular stencils and spread evenly. The electrodes were left at room temperature for 2 hr in a covered glass Petri dish to allow for polystyrene beads to settle and dry. Following this, annealing was performed at 80 °C for 10 min. Meanwhile, 80 mg of ITO nanoparticles (Merck, 544876, <50 nm) were bath sonicated at 37 kHz in 300 µL 5:1 methanol:Milli-Q® for 6 hr. 7.5 µL of the suspension was drop cast evenly onto the polystyrene scaffold and left until it had fully infiltrated the structure. Electrodes were then heated in a Carbolite ELF 11/14B/301 furnace at 500 °C for 20 min from room temperature with a temperature ramp of 1 °C $min^{-1}$ to calcinate the polystyrene beads.

IO-ITO electrodes for spectroelectrochemistry were made by the same protocol with the following modifications. ITO-coated glass (Diamond Coatings, 4.5 Ω $sq^{-1}$) cut into 0.5 cm x 6 cm rectangles was used as a support. 10 µL of a 10% (w/w) solution of polystyrene beads (Merck, 72986, 10 µm) was dropcast into the circular stencils and spread evenly. Electrodes were left at 4 °C overnight followed by 1 hr at room temperature to settle and dry. Annealing was performed at 100 °C for 10 min. 1 µL of ITO nanoparticle suspension was drop cast on the polystyrene scaffold.

#### Biomembrane electrochemistry

##### Electrode modification

Thylakoid membrane modification of electrodes was performed by drop casting thylakoid membrane suspension over the structured area of the electrode in a circular motion (10 µL for photoelectrochemistry and 3.7 µL for spectroelectrochemistry), with care taken to spread the solution evenly across the electrode structure. Electrodes were then left to dry for ~20 min at room temperature, after which they had a glazed (as opposed to wet or cracked) appearance and were used in electrochemistry experiments.

##### Photoelectrochemistry

Photoelectrochemistry of thylakoid membranes was performed in a water-jacketed, glass photoelectrochemical cell with a PTFE lid, maintained at 25 °C and containing 4.5 mL of electrolyte. Measurements were performed with an Ivium Technologies CompactStat.h potentiostat using a thylakoid membrane-modified IO-ITO working electrode, platinum mesh counter electrode and Ag/AgCl reference electrode. Illumination was provided with Thorlabs LED light sources (M680L4, M700L4, M730L5) powered by an LED driver ) and aligned with a collimator. An additional FB730-10 - Ø1" Bandpass Filter filter (2.54 mm, Thorlabs) was used with the 730 nm LED.

##### Spectroelectrochemistry

Spectroelectrochemistry of thylakoid membranes was performed in a photoelectrochemical cell made from a 4 mL quartz crystal cuvette with a PTFE lid, maintained at room temperature and containing 2.4 mL of electrolyte. Measurements were performed with an Ivium pocketSTAT2 potentiostat using a platinum mesh counter electrode and platinum pseudoreference. $E_{app}$ values were calibrated by performing cyclic voltammetry of 500 µM ferricyanide and 500 µM ferrocyanide with a glassy carbon working electrode and comparing the measured $E_m$ with those measured in a glass vial using the same working and counter electrode with a Ag/AgCl reference electrode. Measurements were performed alongside spectroscopic measurements within a JTS-150, which also provided illumination from the 630 nm actinic light ring. These measurements were otherwise performed identically to standard photoelectrochemistry experiments.

##### Standard conditions

Unless otherwise stated, all measurements were performed in following standard conditions: chl *a* loading, 20 µg; illumination, 50 µmol photons $m^{-2}$ $s^{-1}$ of chopped 680 nm light; electrolyte, electrolyte buffer (50 mM MES-NaOH, 15 mM NaCl, 5 mM $MgCl_2$, 2 mM $CaCl_2$, adjusted to pH 6); sampling rate, 10 $s^{-1}$; purging, $N_2$ gas for 20 min before and 5 min after addition of the working electrode with the purging needle moved to the headspace during measurement. Chronoamperometry scans were performed with chopped light (1 min light / 1 min dark) and an applied working electrode potential ($E_{app}$) of +300 mV vs SHE following 5 min pre-equilibration, unless otherwise stated. When possible, three successive photocurrent measurements were performed and average (see below). Stepped chronoamperometry scans were recorded with chopped light (90 s dark / 30 s light / 30 s dark) for measurements at each $E_{app}$ value which ranged -100 to +700 mV vs SHE without, or -600 to +700 mV with, enzymatic oxygen removal provided by 5 mM glucose (Merck, G8270), 1 mg mL$^{-1}$ glucose oxidase (Merck, G7141), and 500 µg mL$^{-1}$ catalase (Merck, C9322)[7]. Measurements were performed following 10 min of pre-equilibration, with two measurements of the first $E_{app}$ value being recorded with only the latter being used in data analysis.

##### Other experimental conditions

Some experiments were performed in non-standard conditions. These include the use of different light conditions (intensity, wavelength or chopping), no purging, different pH, or differently electrolytes such as electrolyte buffer - salts (50 mM MES-NaOH, adjusted to pH 6) and phosphate buffer (43.2 mM $NaH_2PO_4$, 6.8 mM $Na_2HPO_4$, adjusted to pH 6). For some experiments the electrolyte was supplemented with different concentrations of the following chemicals: Carbonyl cyanide m-chlorophenyl hydrazone (CCCP; Merck, C2759; stock, 100 mM in DMSO), 3-(3,4-dichlorophenyl)-1,1-dimethylurea (DCMU; Merck, D2425; working, 100 µM; stock, 100 mM in DMSO), N-oxo-2-heptyl-4-Hydroxyquinoline (HQNO; Cayman Chemical, 15159; working, 10 µM; stock, 1 mM in DMSO), potassium ferricyanide (Merck, 244023; working, 1 mM; stock, 500 mM in electrolyte buffer), $NADP^+$ (Cayman Chemical, 21045; working, 1 mM; stock, 50 mM in electrolyte buffer), NADPH (Merck, 481973; working, 1 mM; stock, 50 mM in electrolyte buffer), PMA (Merck, P27127; stock, 50 mM in DMSO), soluble fraction (working, 160 ng µL$^{-1}$ prior to any filtration), succinate (Merck, 14160; working 5 mM; stock, 1M in electrolyte buffer). Chemical titrations were performed by successively adding chemical stocks to the electrolyte between measurements. The electrolyte was mixed with a 5000 µL pipette, with 5 min of purging prior to each measurement. pH titrations were performed from high to low pH by replacing the electrolyte with a new solution adjusted to the desired pH with HCl and NaOH, with 20 min of purging prior to each measurement. Photocurrent measurements with different light intensities were performed by increasing the power of the LED in the dark period between photocurrent measurements. Photocurrent measurements with different dark adaptation times were changing the programmed light chopping in the dark period between photocurrent measurements.

#### Photocurrent data analysis

Analysis and plotting of thylakoid membrane photocurrents were performed in bespoke software developed in MATLAB. The software accepts raw data files as inputs, alongside various experimental and other parameters. An option is included to normalize the data by chl content (in nmol), which is provided by the user. Analysis of chronoamperometry and stepped chronoamperometry data was performed by processing the data from individual photocurrent measurements in steps depicted in Extended Data Fig. 4b, with example code provided (Supplementary Code).

##### Dark current measurement

Dark currents were calculated from raw photocurrent data as follows:

$$I_{dc}\ (\text{A}) = \frac{\sum_{t=t_{\text{deq}}}^{t_{\text{end}}} I_{\text{exp}}(t)}{t_{\text{end}} - t_{\text{deq}}}$$

where $I_{dc}$ is the dark current, $I_{\text{exp}}(t)$ is the experimental current data at time $t$, $t_{\text{deq}}$ is the dark equilibration time (time at which the dark current reaches steady state) and $t_{\text{end}}$ is the final timepoint of the data.

##### Baselining

Baselining was performed by fitting to the dark signal. Fitting was performed with a straight line for most data. For electrochemical data in which a straight line provided a poor fit (especially true of stepped chronoamperometry data), dark current was fitted with the following treatment of the Cottrell equation for a spherical electrode[8]:

$$I_{\text{fit}}\big(t \mid 0 \le t < t_{on} \cup t_{\text{deq}} < t \le t_{end}\big)\ (\text{A}) = nFA\left(\frac{\sqrt{Dc_0}}{\sqrt{\pi t}} + \frac{Dc_0}{r}\right)$$

where $I_{\text{fit}}(t)$ is the fitted current at time $t$, $t_{on}$ is the time at the start of the light period, $t_{\text{deq}}$ is the dark equilibration time (time at which the dark current reaches steady state), $t_{\text{end}}$ is the final timepoint of the data, $n$ is the number of electrons involved in the redox reaction of the analyte, $F$ is Faraday's constant, $A$ is the planar surface area of the electrode, $D$ is the diffusion coefficient of the analyte, $c_0$ is the concentration of analyte at $t = 0$, and $r$ is the radius of the electrode. All parameters were fitted apart from $r$, which was set as 5 mm. Fitting was performed using the fminsearch function to minimize the sum of squared estimate of errors ($SSE$) between the data and the fit, as follow:

$$SSE = \sum_{t=0}^{t_{\text{end}}} \left[I_{\text{exp}}(t) - I_{\text{fit}}(t)\right]$$

where $I_{\text{exp}}(t)$ and $I_{\text{fit}}(t)$ are the experimental and fitted current at time $t$, and $t_{\text{end}}$ is the final timepoint of the data. Because the position of $t = 0$ is not known, curve fitting is performed at increasing values of the parameter until an optimal fit is identified.

In stepped chronoamperometry experiments the dark current was found to be offset between different replicates, likely caused by differences in electrode capacitance at more positive $E_{\text{app}}$ values. To account for this, dark currents were shifted to an average value prior to baselining, as follows:

$$I_{\text{adj}}(n,t)\ (\text{A}) = I_{\text{exp}}(n,t) - \left(I_{\text{exp}}(n, t_{\text{end}}) - \frac{\sum_1^N I_{\text{exp}}(n, t_{\text{end}})}{N}\right)$$

where $I_{\text{exp}}(n,t)$ $I_{adj}(n,t)$ are the experimental and adjusted experimental current for replicate $n$ at time $t$, $t_{\text{end}}$ is the final timepoint of the data, and $N$ is the total number of replicates.

Final fitted curves were subtracted from experimental data to provide baselined photocurrent data:

$$I_{\text{base}}(t)\ (\text{A}) = I_{\text{exp}}(t) - I_{\text{fit}}(t)$$

where $I_{\text{exp}}(t)$ and $I_{\text{base}}(t)$ are the experimental and baselined current at time $t$.

#### Photocurrent parameter measurements

Steady State Photocurrents were calculated from baselined photocurrents as follows:

$$I_{ssp}\ (\text{A}) = \frac{\sum_{t=t_{\text{leq}}}^{t_{\text{off}}} I_{\text{base}}(t)}{t_{\text{off}} - t_{\text{leq}}}$$

where $I_{ssp}$ is the Steady State Photocurrent, $t_{\text{off}}$ is the time at the end of the light period and $t_{\text{leq}}$ is the light equilibration time (time at which the light current reaches steady state), and $I_{\text{base}}(t)$ is the baselined current at time $t$ .

Spike Charges were calculated from baselined photocurrents using the trapz function as follows:

$$Q_{sc}\ (\text{C}) = \int_{t_{\text{on}}}^{t_{\text{off}}} \left[I_{base}(t) - I_{ssp}\right] dt$$

where $Q_{sc}$ is the Spike Charge, $t_{\text{on}}$ and $t_{\text{off}}$ are the times at the start and end of the light period, $I_{\text{base}}(t)$ is the baselined current at time $t$, and $I_{ssp}$ is the Steady State Photocurrent.

#### Averaging replicates

For chronoamperometry data, three successive photocurrents were recorded for each tested condition. The mean of these photocurrents and their respective parameters were calculated to reduce noise. For all photocurrent measurements, the mean and standard error of the mean (S.E.M.) were calculated for photocurrents and their parameters from three independent biological replicates.

#### Cyclic voltammetry

All cyclic voltammetry experiments were performed with an Ivium Technologies CompactStat.h following 20 minutes of purging with N2 gas, with the purging needle being moved to the headspace of the electrochemical cell during measurement.

Cyclic voltammetry of lysis buffer, soluble fractions and the ferri-/ferrocyanide redox couple (500 μM ferricyanide and 500 μM ferrocyanide) was performed in a 20 mL glass filled with 5 mL of solution, using an ITO-coated glass working electrode, platinum mesh counter electrode, and Ag/AgCl reference electrode. Measurements were performed following a 1 min pre-equilibration at -600 mV vs SHE, after which $E_{app}$ was changed from -600 to +700 mV vs SHE with a scan rate of 10 mV $s^{-1}$. The third and final scan was plotted. Soluble fractions were not diluted prior to measurement.

All other cyclic voltammetry experiments were performed with the same electrochemical apparatus used in thylakoid membrane photoelectrochemistry experiments. Measurements were performed following a 5 min pre-equilibration at +800 mV vs SHE, after which $E_{app}$ was changed from +800 to -200 mV vs SHE with a scan rate of 2 mV $s^{-1}$. The first and only scan was plotted. Measurements of different electrolytes and redox molecules were performed with blank IO-ITO electrodes. Measurements of redox molecules were performed in electrolyte buffer with the following concentrations: 10 μM HQNO, 1 mM ferri-/ferrocyanide redox couple (500 μM ferricyanide and 500 μM ferrocyanide), 500 μM $NADP^+$ and 500 μM NADPH. Measurements of thylakoid membrane-modified IO-ITO electrodes were performed in darkness or under 50 μmol photons $m^{-2}$ $s^{-1}$ of 680 nm in the presence of either 1% DMSO, 100 μM DCMU or 10 μM HQNO.

The position of redox peaks was determined using the "findpeaks" function in MATLAB when possible, with other clearly visible peaks being annotated manually. $E_m$ values were calculated as the midpoint between a pair of oxidation and reduction peaks.

#### Structural and chemical analysis of isolated membranes

##### Scanning transmission electron microscopy

Scanning transmission electron microscopy of isolated thylakoids membranes was performed with a TESCAN MIRA3 FEG-SEM with beam acceleration of 30 kV. Samples were diluted to a chl *a* concentration of 1 ng $\mu L^{-1}$ in deionized water. 2 μL of this solution was pipetted onto a holey carbon-supported copper TEM grid (200 mesh) and left to dry at 4 °C for 2 hr prior to imaging.

##### Nanoparticle tracking analysis

Nanoparticle tracking analysis was performed using a Malvern Panalytical Nanosight NS500 fitted with an Electron Multiplying Charge Coupled Device camera configured with a 522 nm laser. Particle concentrations were determined by tracking and counting particles in microscopy images. Particle sizes were determined based on the diffusion speed of tracked particles. Thylakoid membrane samples were diluted to a chl *a* concentration of 10 pg $\mu L^{-1}$ in electrolyte buffer, with 5 x 60 s videos recorded for each sample and analyzed with NTA 3.2 software using a detection threshold of 5.

##### Membrane topology analysis

The polarity of cyanobacteria TMs was determined by 2-phase polymer partitioning[9]. All steps were performed at 4 °C. Solutions of dextran-500 (11.6% w/w) and PEG-3350 (11.6% w/w) were prepared in partitioning buffer (5 mM K2HPO4, 250 mM sucrose, adjusted to pH 7.8 with HCl or NaOH). 20 μg of isolated thylakoid membranes were deposited into a 2 mL microcentrifuge tube containing 750 μL of PEG-3350 solution layered on top of 750 μL of dextran-500 solution. Tubes were mixed by inverting 35 times before phase partitioning by centrifugation (1000 x g, 4 min). The upper phase was added to a fresh 2 mL microcentrifuge tube containing 750 μL of dextran-500 partitioning buffer. The lower phase was topped with 750 μL of fresh PEG-3350 partitioning buffer. Mixing and phase partitioning was subsequently repeated twice, after which the upper and lower phases from all tubes were pooled into separate 15 mL centrifuge tubes and subjected to one final phase partitioning. The upper and lower phases from each tube were again pooled in 50 mL centrifuge tubes, diluted up to 30 mL with partitioning buffer, and centrifuged (50,000 x g, 15 min) to pellet the thylakoid membranes. Supernatants were discarded, with pellets being resuspended in 1 mL of methanol and transferred to 1.5 mL microcentrifuge tubes. Centrifugation (13,000 x g, 3 min) was used to remove debris, after which the chl *a* concentrations of the supernatants were determined. The proportion of thylakoids which were cytoplasmic side-out was determined as follows:

$$\text{Cytoplasmic side–out} = \frac{[\text{Chl a}]_{\text{lower phase}}}{[\text{Chl a}]_{\text{lower phase}} + [\text{Chl a}]_{\text{upper phase}}}$$

#### Fluorimetry

Excitation-emission matrices were recorded with an Agilent Cary Eclipse Fluorescence Spectrometer. Measurements were performed with excitation and emission wavelengths ranging from 200-800 nm, using a step size of 5 nm, scan rate of 1000 nm $min^{-1}$, averaging time of 0.3 s, excitation and emission bandwidths of 5 nm, and a PMT detector sensitivity of 500 V. For measurements of isolated thylakoid membranes, a chl *a* concentration of 10 ng $\mu L^{-1}$ was used. For measurements of soluble fractions, a (pre-filtration) C-phycocyanin concentration of 160 ng $mL^{-1}$ was used.

#### Pigment quantification

UV-visible spectroscopy was performed using a Shimadzu UV-1800 Spectrophotometer. Absorbance spectra were recorded in plastic 1 mL cuvettes from 350 to 800 nm using a suitable blank.

Chl *a* concentrations of cyanobacterial cultures were calculated as follows[10]:

$$[\text{Chl } a]\ (\mu\text{g}\ \mu\text{L}^{-1}) = \frac{\text{OD}_{680} - \text{OD}_{750}}{103.1}$$

Chl *a* concentrations of cyanobacterial thylakoid membrane samples were determined in methanol as follows[11]:

$$[\text{Chl } a]\ (\mu\text{g}\ \mu\text{L}^{-1}) = \frac{\text{OD}_{665} - \text{OD}_{750}}{79.95}$$

Chl *a* and *b* concentrations of algal and plant thylakoid membrane were determined in a 4:1 acetone to methanol mixture as follows[11]:

$$[\text{Chl } a + b]\ (\mu\text{g}\ \mu\text{L}^{-1}) = \frac{17.76(\text{OD}_{647} - \text{OD}_{750}) + 7.34(\text{OD}_{664} - \text{OD}_{750})}{1000}$$

The C-phycocyanin concentration of the soluble fraction diluted in lysis buffer was determined as follows[12]:

$$[\text{C-phycocyanin}]\ (\mu\text{g}\ \mu\text{L}^{-1}) = \frac{\text{OD}_{620} - (0.7 \times \text{OD}_{750})}{7.38}$$

#### Protein gel electrophoresis

BN-PAGE was performed using isolated thylakoid membranes diluted to a chl *a* concentration of 1 mg $mL^{-1}$. 5 µL of this solution was mixed with 5 µL of resuspension buffer (1M Bis-Tris HCl, 50% glycerol, 10 mg $mL^{-1}$ Pefabloc, 1M sodium fluoride, 2% n-decyl-beta-maltoside, adjusted to pH 7) by gently pipetting up and down on ice. The solution was centrifuged at 4 °C (18,000 x g, 20 min), after which the supernatant was loaded onto the gel (Invitrogen™ NativePAGE™ Bis-Tris Mini Protein Gels, 4 to 16%, 1.0 mm). The gel was run with separate running buffers for the anode (50 mM Bis-Tris/HCl, adjusted to pH 7) and cathode (15 mM Bis-Tris/HCl, 50 mM Tricine, 0.01% Serva Blue G, adjusted to pH 7) at the following voltages: 75 V for 30 min, 100 V for 30 min, 125 V for 30 min. The cathode buffer was subsequently removed, and replaced with fresh cathode buffer without any Serva Blue G, after which the gel was ran to completion at the following voltages: 150 V for 60 min, 175 V 30 min.

SDS-PAGE was performed with cyanobacterial lysate, soluble fraction, and isolated thylakoid membranes. Total protein concentrations of each sample were determined by a Bradford assay against a standard curve obtained with 0, 10, 50, 250, 500 and 1000 µg $mL^{-1}$ Bovine Serum Albumin (Merck, A2153), with 30 µg of protein in each lane. Samples were mixed with an appropriate volume of 4X Laemmli buffer in PCR tubes and heated to 100 °C for 10 min. Samples were then centrifuged (13,000 x g, 5 min) with the resulting supernatant loaded onto a BioRad Mini-PROTEAN® TGX® Stain-Free Gel (4-20%). 7 µL of PageRulerTM Plus Prestained Protein Ladder (10 to 250 kDa) was used. Gels were run for 75 min at 100 V in SDS running buffer (25.01 mM Tris base, 192.4 mM glycine, 3.467 mM SDS).

#### Measurements of photosynthetic activity

##### Oxygen evolution measurements

Oxygen evolution measurements were recorded with a Pyroscience FireSting®-PRO using a OXROB10-HS High Speed Robust Oxygen Probe inserted into 1 mL of solution within a 5 mL glass vial. Thylakoid membrane suspensions were prepared by diluting isolated thylakoid membranes in electrolyte buffer (pH 6). Cell solutions were prepared by concentrating cells in BG-11 medium (pH 7.8). Measurements were performed at 25 °C with stirring at 900 rpm. PSII oxygen evolution measurements were performed with a chl *a* concentration of 40 µg $mL^{-1}$ in the presence of 1 mM DCBQ (Merck, 431982) and 1 mM potassium ferricyanide as electron acceptors[3]. PSI oxygen evolution measurements were performed with a chl *a* concentration of 20 µg $mL^{-1}$ in the presence of 100 µM DCMU, 20 mM ascorbate (Merck, 11140) and 200 µM DCPIP (Merck, D1878) as electron donors, with 4 mM methyl viologen (Merck, 856177) as an electron acceptor[13]. Samples were left to pre-equilibrate to a steady $O_2$ level prior to measurement under 1500 µmol photons $m^{-2}$ $s^{-1}$ of chopped 680 nm (2 min light, 2 min dark). Data was baselined to the pre-equilibration signal and subsequently analyzed by independently fitting the light and dark currents to the following equation:

$$[O_2]\ (\mu\text{mol L}^{-1}) = (k_{O_2} \times t) + \text{c}$$

where $k_{O_2}$ is the $O_2$ production rate (µmol $s^{-1}$), $t$ is time (s) from beginning of the light/dark period and c is the $[O_2]$ at $t$=0. The electron transfer rate, measured as the absolute oxygen evolution rate, was calculated as follows:

$$\text{Electron Transfer Rate } (\Delta\text{nmol O}_2\ [\text{nmol chl a}]^{-1}\ \text{hr}^{-1}) = \left|\frac{3600\left(k_{O_2}^{light} - k_{O_2}^{dark}\right)}{\text{Chl a}}\right|$$

where $k_{O_2}^{light}$ and $k_{O_2}^{dark}$ are the $O_2$ production rates (µmol $s^{-1}$) in the light and dark, with the mass of Chl *a* expressed in nmol.

##### Chlorophyll fluorescence

Chl fluorescence measurements were performed using a BioLogic JTS-150 Spectrometer at room temperature using a 4 mL crystal quartz cuvette containing 2 mL of electrolyte buffer supplemented with cyanobacterial thylakoid membranes to a final chl *a* concentration of 10 µg $mL^{-1}$. Measurements were performed under increasing intensities of chopped 630 nm light (1 min light / 1 min dark) using 1 mM potassium ferricyanide as an electron acceptor. A 450 nm measuring light was used as an excitation beam, with a 700 nm High Performance long-pass filter (5 mm, Edmund Optics) on the measuring detector and a BG39 filter (3 mm, Schott) on the reference detector. A detector current of 50 mA was used with 150 µs detecting pulses recorded within a 1 ms dark interval. The sample was further excited with a 99 ms multiple turnover flash of 7000 µmol $m^{-2}$ $s^{-1}$ at 40 s into the light and dark period. PSII quantum efficiency was calculated as follows[14]:

$$\Phi_{\text{PSII}} = \frac{F_{\text{m}}' - F_{\text{m}}}{F_{\text{m}}'}$$

where $\Phi_{\text{PSII}}$ is the PSII quantum efficiency, and $F_{\text{m}}$ and $F_{\text{m}}'$ are the maximum fluorescence in the dark and light respectively.

#### Redox cofactor absorbance

Redox cofactor absorbance measurements were performed using a BioLogic JTS-150 Spectrometer at room temperature using a 4 mL crystal quartz cuvette containing 2 mL of electrolyte buffer supplemented with cyanobacterial thylakoid membranes to a final chl *a* concentration of 10 µg $mL^{-1}$.

Cytochrome *b* and *f* redox changes were recorded under increasing intensities of chopped 630 nm light (1 min light / 1 min dark) using 1 mM potassium ferricyanide as an electron acceptor. 546, 554, 563 and 574 nm measuring lights were used, with BG39 filters (3 mm, Schott) on the measuring and reference detectors. A detector voltage of 5V was used with 150 µs detecting pulses recorded within a 1 ms dark interval. The amount of oxidized cytochrome *b* and *f* were calculated as follows[15]:

$$[\text{Cyt } b_{\text{ox}}]_{\text{exp}}(t)\ \left(\frac{\Delta \text{I}}{\text{I}}\right) = A_{563}(t) - (0.607A_{574}(t) + 0.393A_{546}(t))$$

$$[\text{Cyt } f_{\text{ox}}]_{\text{exp}}(t)\ \left(\frac{\Delta \text{I}}{\text{I}}\right) = A_{554}(t) - (0.607A_{574}(t) + 0.393A_{546}(t))$$

where $[\text{X}_{\text{ox}}](t)$ is the experimental amount of oxidized cofactor and $A_x(t)$ is the absorbance change recorded at wavelength $x$ and time $t$.

Plastocyanin and P700 redox changes were recorded under increasing intensities of chopped 630 nm light (1 min light / 1 min dark) using 1 mM methyl viologen as an electron acceptor. 705 and 740 nm measuring lights were used, with RG695 filters (3 mm, Schott) on the measuring and reference detectors. A detector voltage of 4V was used with 100 µs detecting pulses recorded within a 1 ms dark interval. The amount of oxidized plastocyanin and P700 were calculated as follows[16]:

$$[\text{Plastocyanin}_{\text{ox}}]_{\text{exp}}(t)\ \left(\frac{\Delta \text{I}}{\text{I}}\right) = A_{740}(t)$$

$$[\text{P700}^{+}]_{\text{exp}}(t)\left(\frac{\Delta \text{I}}{\text{I}}\right) = A_{705}(t) - A_{740}(t)$$

where $[\text{X}](t)$ is the experimental amount of oxidized cofactor and $A_x(t)$ is the absorbance change recorded at wavelength $x$ and time $t$.

Photoresponses were baselined to the dark signal (see above). Steady state redox changes of cofactors were calculated from baselined redox changes as follows:

$$[\text{X}]_{ss}\ \left(\frac{\Delta \text{I}}{\text{I}}\right) = \left|\frac{\sum_{t=t_{\text{leq}}}^{t_{\text{off}}}[\text{X}]_{\text{base}}(t)}{t_{\text{off}} - t_{\text{leq}}}\right|$$

where $[X]_{ss}$ is the steady state redox change of cofactor X, $t_{\text{leq}}$ is the light equilibration time (time at which the light current reaches steady state), $t_{\text{off}}$ is the time at the end of the light period, and $[\text{X}]_{\text{base}}$ is the baselined amount of oxidized cofactor at time $t$. The absolute value of this parameter corresponds to the oxidation of P700 and cytochromes *b* and *f*, and the reduction of plastocyanin.

The rate of P700 re-reduction in the dark was calculated by fitting the following exponential decay curve to the first 7 seconds of the dark signal[17]:

$$[\text{P700}^{+}]_{\text{base}}(t \mid t_{\text{off}} \leq t < t_{\text{off}} + 7)\ \left(\frac{\Delta \text{I}}{\text{I}}\right) = Be^{-t/\tau} + [\text{P700}^{+}]_{\text{base}}(t_{\text{off}})$$

where $[\text{P700}^{+}]_{\text{base}}(t)$ is the baselined amount of oxidized P700, $t_{\text{off}}$ is the time at the end of the light period, $B$ is a scaling factor, and $\tau$ is the time constant. The rate of P700 re-reduction was subsequently calculated as the inverse of $\tau$.

#### NADPH fluorescence

NADPH fluorescence measurements were performed using a BioLogic JTS-150 Spectrometer at room temperature using a 4 mL crystal quartz cuvette containing 2 mL of electrolyte buffer supplemented with cyanobacterial thylakoid membranes to a final chl *a* concentration of 10 µg $mL^{-1}$. Measurements were performed under 500 µmol $m^{-2}$ $s^{-1}$ of chopped 630 nm light (1 min light / 10 min dark) using 1 mM $NADP^{+}$ as an electron acceptor. A 385 nm measuring light was used as an excitation beam, with a BG39 filter (3 mm, Schott) and a custom 500 nm long-pass filter on the measuring detector, with a BG39 filter (3 mm, Schott) on the reference detector[18]. A detector current of 100 mA was used with 100 µs detecting pulses recorded within a 1 ms dark interval. Photoresponses were baselined to the dark signal (see above).

#### Electrochromic shift

Electrochromic shift measurements were performed using a BioLogic JTS-150 Spectrometer at room temperature using a 4 mL crystal quartz cuvette containing 2 mL of electrolyte buffer supplemented with cyanobacterial thylakoid membranes to a final chl *a* concentration of 10 µg $mL^{-1}$. Measurements were performed under 500 µmol photons $m^{-2}$ $s^{-1}$ of 630 nm light (1 min light / 1 min dark) using 1 mM potassium ferricyanide as an electron acceptor. 480 and 500 nm measuring lights were used[18], with BG39 filters (3 mm, Schott) on the measuring and reference detectors. A detector voltage of 5 V was used with 100 µs detecting pulses, recorded within a 200 µs dark interval for the final 10 s of the light period and first 10 s of the dark period, or within a 1 ms dark interval otherwise. Photoresponses were baselined to the dark signal (see above).

### Characterisation of the bio-electrode interface:

#### Scanning electron microscopy

Scanning electron microscopy of electrodes was performed with a TESCAN MIRA3 FEG-SEM with beam acceleration of 30 kV. Where necessary, samples were left to dry at 4 °C overnight. Prior to imaging, samples were sputter coated with a 10 nm Pt layer using a Quorum Technologies Q150T ES Turbo-Pumped Sputter Coater.

#### Quantum crystal microbalance with dissipation

Quartz Crystal Microbalance with Dissipation (QCM-D) was performed using a Biolin Scientific QSense E4 QCM with gold sensors (Biolin Scientific, QSX 301) sputter coated with a 10 nm layer of ITO using a Quorum Technologies Q150T ES Turbo-Pumped Sputter Coater. Sensors were cleaned prior to use by being submerged in 1% Hellmanex II for 30 min, rinsed with ultrapure water, and dried with $N_2$ gas. Sensors were then sonicated in 99% ethanol for 10 min, rinsed with ultrapure water, dried with $N_2$ gas, and UV-ozone treated for 10 min using a Bioforce Nanosciences UV-Ozone Procleaner. A flow rate of 150 µL $min^{-1}$ was used for all experiments. After each solution was pumped into the flow cell the pumping was stopped. The sensors were left until no observable changes in frequency or dissipation were observed before further solutions were pumped in. Sensors were pre-equilibrated in electrolyte buffer for 1 hr before measurements. Thylakoid membrane suspensions were diluted to a chl *a* concentration of 100 ng $\mu L^{-1}$ in storage buffer prior to being pumped into the flow cell.

#### Protein structure analysis

Structure of photosynthetic complexes were visualized with UCSF ChimeraX[19]. Models of PSII dimers (isolated from *Thermostichus vulcanus*, PDB: 3WU2[20]), cytochrome b6f dimers (isolated from *Nostoc* sp. PCC 7120, PDB: 4H44[21]), and PSI trimers (isolated from *Thermosynechococcus vestitus* BP-1, PDB: 1JB0[22]) embedded within simulated lipid bilayers were obtained from the MemProtMD database[23]. Ligands were added to models by alignment with the biological assemblies obtained from the PDB database. In the case of PSI, molecular dynamics simulations had only been performed on the protein monomer, despite the biological assembly being a trimer. A lipid bilayer model for the trimer was made by aligning the MemProtMD structure with each of the monomer chains, followed by the deletion of any clashing lipid residues (van der Waals overlap ≥ 0.6 Å after subtracting 0.4 Å for hydrogen bonding).

#### Supporting Results 1: Characterisation of isolated thylakoid membranes

Previous methods for crude cyanobacterial thylakoid membrane extraction[3] were adapted to isolate enriched thylakoid membranes from the model cyanobacterium *Synechocystis* sp. PCC 6803 (Figure S1). Based on chlorophyll (chl) *a* concentration, this isolation method enabled ~33% of the thylakoid membrane to be recovered, yielding vesicles with a diameter of 100-250 nm, as measured by electron microscopy (Figure 1e) and nanoparticle tracking analysis (Figure S2a), with polymer 2-phase separation[9] revealing a preferential orientation with the cytoplasmic face outwards in 84.3±2.5% of vesicles. Maintaining this topology is essential to prevent abiotic and competing electron transfer reactions[7].

The presence of certain thylakoid membrane proteins can be determined by the spectroscopic signatures of their bound pigments. This includes the chl *a* bound to photosystems and the phycobilins bound to the light harvesting phycobilisomes. Isolated thylakoid membrane samples exhibited clear absorbance and fluorescence signatures for chl *a* but lacked those for phycobilisomes, many of which detached from the membranes during extraction (Figure S2b-c). However, SDS-PAGE analysis showed the presence of bands characteristic of the phycobiliproteins CpcA/B in thylakoid membrane samples (suggesting some phycobilisomes remain attached), alongside other bands corresponding to thylakoid membrane proteins[24] (Figure S3a). BN-PAGE demonstrated the presence and integrity of major photosynthetic and respiratory thylakoid membrane complexes in the samples (Figure S3b), consistent with results from previous studies[25].

The ability of the thylakoid isolation method (Figure S1) to obtain photosynthetically active samples was tested by measuring the maximal rates of PSII and PSI electron transfer in oxygen evolution experiments[3,13]. Activity of both photosystems was demonstrated (Figure S4a), with the enhanced rate of PSI being consistent with its increased and decreased electron transfer lifetime.

A suite of spectroscopic methods were subsequently used to determine the functioning of the entire thylakoid membrane photosynthetic electron transport chain[26] (Fig. 2a) under physiologically-relevant conditions. Photosystem II (PSII) activity was determined through measurements of its quantum efficiency ($\Phi_{PSII}$)[14], which were consistently >0.4 at all measured light intensities (Figure S4b). Cytochrome $b_6f$ redox changes were also measured[15], with maximal oxidation of both *b* and *f* cytochromes being achieved at a light intensity of 500 μmol photons $m^{-2}$ $s^{-1}$ (Figure S4c). The PSII $Q_B$ inhibitor DCMU[27] inhibited both $\Phi_{PSII}$ and cytochrome oxidation, consistent with these processes being dependent on photocatalytic water oxidation by PSII.

Reduction of plastocyanin and oxidation of the P700 primary donor of Photosystem I (PSI)[16] were also observed upon illumination (Figure S4d-e), demonstrating photosynthetic electron transport activity. These redox changes were insensitive to both light intensity and DCMU addition, consistent with PSI being donor-side limited, as is often the case *in vivo*. However, it is important to note that measurements of plastocyanin and P700 redox changes were recorded using methyl viologen as an electron acceptor which readily reacts with oxygen to generate reactive oxygen species, thereby leading to protein damage which will diminish redox signals[28].

NADPH fluorescence of thylakoid membrane samples[29] was also measured in the absence of exogenous $NADP^+$ addition, with no clear NADPH synthesis being observed upon illumination with 500 μmol photons $m^{-2}$ $s^{-1}$ (Figure S4f). This result demonstrates the absence of ferredoxin and ferredoxin-$NADP^+$ reductase (FNR) in isolated thylakoid membrane samples. However, an increase in NADPH fluorescence was observed in the extended dark period, suggesting the presence of a light-independent NADPH synthesis pathway. DCMU addition had no effect on this light-independent pathway.

Electrochromic shift measurements were used to measure the strength of the proton motive force across the thylakoid membrane[30,31]. No decay in the ECS signal was observed at light-dark transition (Figure S4g), consistent with there being no proton motive force formed across the thylakoid membrane during illumination. This can be attributed to the isolated thylakoid membranes being permeable to protons, either through rapid activity of the ATP synthase and/or due to ion leakage occurring across the membrane.

Taken together, these results demonstrate light-dependent activity of the entire photosynthetic electron transport chain, from PSII to FNR, with no inhibition of the electron transfer rate during long-term illumination caused by the formation of a proton motive force.

#### Supporting Results 2: Membrane-modification of electrodes

To enable high loading and efficient wiring of isolated thylakoid membranes to electrodes, hierarchically structured and translucent inverse opal-indium tin oxide (IO-ITO) electrodes[6] were fabricated with a pore diameter of 10 µm and solvent channel diameter of 2 µm (Figure S5), enabling thylakoid membrane vesicles to penetrate the electrode structure. Thylakoid membrane-modified IO-ITO electrodes were created by dropcasting a thylakoid membrane suspension onto a bare electrode, leading to the formation of a multi-membrane thick layer coating the ITO-nanoparticles throughout the electrode structure (Fig. 1c-d and S6). QCM-D measurements of thylakoid membrane adsorption to ITO-coated chips revealed a large monophasic decrease in frequency and increase in dissipation (Figure S7), consistent with the adsorption of vesicles to the ITO[32].

#### Supporting Results 3: Analysis of thylakoid membrane photoelectrochemistry data

A standard pipeline was developed to enable reproducible analysis of thylakoid membrane photoelectrochemistry data (Figure S9). Software was developed in MATLAB to perform the analysis pipeline (Materials and Methods).

Chronoamperometry scans of thylakoid membrane-modified electrodes were performed under chopped illumination, giving rise to successive photocurrent profile measurements (Figure S9). Scans were performed until photocurrent profiles stabilised, which generally occurred by the second measured profile. Data from the first photocurrent profile was excluded to ensure the dark adaptation time was consistent for all measurements. The analysis pipeline begins by segmenting the current data into individual photocurrent profile measurements with each consisting of the current measured i) shortly before illumination, ii) during the entire light periods, iii) during the entire following dark period (with 5/60/60 second duration for standard experiments). For each photocurrent profile the dark current was recorded (Step ①). After this, baselines were fitted to the dark periods of each photocurrent profile (Step ②). Next, the photocurrent parameters (Steady State Photocurrent and Spike Charge; Fig. 1f) were calculated for each photocurrent profile (Step ③). The effect of electrical noise was diminished by averaging the data from three successive photocurrent profile measurements (Step ④). Finally, the data from photocurrent profiles recorded with multiple biological replicates (generally 3) was averaged, with the error calculated as the standard error of the mean (Step ⑤). Baseline-subtracted photocurrent profiles and measured photocurrent parameters could then be plotted.

For stepped chronoamperometry scans, a single photocurrent profile was recorded at increasing $E_{app}$ values. Data analysis was therefore performed identically as for chronoamperometry scans, albeit with the omission of Step ④ (Figure S9).

#### Supporting Results 4: Effect of experimental conditions on thylakoid membrane photocurrents

Thylakoid membrane photocurrents were recorded across a range of experimental conditions to provide insights into the molecular mechanisms underpinning photocurrent generation, as well as to identify standardized conditions to be used in further photoelectrochemistry experiments.

As expected, increased loadings of thylakoid membranes onto electrodes yielded enhancements in both the Spike Charge and Steady State Photocurrent (Figure S10a) albeit at different rates, with Steady State Photocurrents plateauing with 10 µg of chl *a* and Spike Charges continuing to increase at 20 µg of chl *a*. This is consistent with the Steady State Photocurrents being caused by direct electron transfer to the electrode, which reaches its maximum when the electrode surface is completely covered with thylakoid membranes (Figure S6); with Spike Charges instead stemming from currents (either non-Faradaic and/or Faradaic) caused by mobile charged species trapped within the isolated thylakoid membrane samples (Supporting Results 5). Recording photocurrents under increasing intensities of 680 nm light showed a clear light-dependent enhancement in the Steady State Photocurrent, which was not observed for the Spike Charge (Figure S10b), demonstrating the two features dependent on different electron transport pathways.

Varying the pH revealed an inhibition of the Steady State Photocurrent at more acidic conditions (Figure S10c) with no anodic Steady State Photocurrent detected at pH 4. This can be attributed to inhibition of PSII and Cytochrome $b_6f$ at this pH[33,34]. The Spike Charge showed a less clear pH-dependent relationship, although the spike feature was clearly present at pH 4. The formation of ΔpH across the thylakoid membrane during photosynthesis is known to inhibit the electron transport rate[35]. To test the effect of ΔpH, photocurrents were recorded in the presence of the protonophore carbonyl cyanide m-chlorophenyl hydrazone (CCCP), which dissipates ΔpH[36]. No enhancement of photocurrent features was observed in the presence of CCCP (Figure S10d), suggesting its addition did not lead to enhancement of the electron transfer rate. This result is consistent with the lack of ΔpH measured across the thylakoid membrane in spectroscopic measurements (Supporting Results 1, Figure S4g). An inhibition of the Steady State Photocurrent was observed upon CCCP addition, which is consistent with its known off-target inhibition of PSII water oxidation[37].

The presence and relative magnitudes of the Steady State Photocurrent and Spike Charge at different working electrode potentials were consistent across different electrolytes (Figure S10e), demonstrating that the features were not caused by abiotic interactions with the buffer conditions. The effect of $O_2$ on photocurrents was determined by recording photocurrents with and without purging with $N_2$ gas. The presence of $O_2$ was found to cause an insignificant decrease in both the Steady State Photocurrent and the Spike Charge (Figure S10f), although it should be noted that the purging methodology (Materials and Methods) was probably unable to remove efficiently any $O_2$ generated at the thylakoid membrane by PSII water oxidation[7].

A chl *a* loading of 20 µg (to improve Spike Charge measurements), 680 nm light (equally excites PSII and PSI) at 50 µmol photons $m^{-2}$ $s^{-1}$ (similar to the growth light intensity), pH 6 (lying between the *in vivo* cytoplasmic and lumenal pH in light), purging to remove $O_2$ (to limit photodamage) and use of an applied working electrode potential ($E_{app}$) of +300 mV vs SHE (enabling anodic electron transfer from PSII, Cytochrome $b_6f$ and PSI) were adopted as standard conditions for further experiments (Materials and Methods). Photodegradation of thylakoid membranes under these standard conditions was determined by recording the change in Steady State Photocurrent over time, with ~40% activity being retained after 400 minutes (Figure S10g). This decay in Steady State Photocurrent was slower than that previously measured with isolated PSII[6] and plant thylakoids[38] and ensured that (with appropriate controls) analytical measurements could be performed for up to 2 hours, at which point >80% of Steady State Photocurrent magnitude was maintained.

#### Supporting Results 5: Evidence for direct and mediated electron transfer between thylakoid membranes and the electrode

According to Marcus theory and experimental measurements, redox cofactors can be assumed capable of performing DET to the electrode if they lie within 2 nm of the cytoplasmic face of the thylakoid membrane which contacts ITO[39]. The structures of the major photosynthetic complexes in simulated lipid bilayers[23] (Figure S13) suggest that multiple redox cofactors lie within this range: $Q_A$ and $Q_B$ of PSII; haem $c_n$ of cytochrome $b_6f$; PhQ, $Fe_X$, $Fe_A$ and $Fe_B$ of PSI. Additionally, previous publications have shown the ability of an electrode to oxidize $PQH_2$ within a lipid bilayer directly, albeit at a large overpotential[40]. This suggests the electron transfer between PETC cofactors and the electrode could be dependent on DET.

To test if diffusible biomolecules could facilitate mediated electron transfer between the PETC and the electrode, photocurrents were recorded in electrolyte exogenously supplemented with the soluble fraction obtained during thylakoid membrane isolation (Figure S1). This soluble fraction contained redox proteins and metabolites expected to have been depleted during thylakoid membrane isolation, especially those located within the cytoplasm such as ferredoxin and NADP(H)[26].

Soluble fraction addition led to the $E_{app}$ at which anodic photocurrents appeared being shifted down to -400 mV vs SHE (Figure S14a-b), suggesting that its addition stimulated PSI-dependent electron transfer to the electrode. Anodic photocurrents were also observed at lower $E_{app}$ values with soluble fraction passed through a 3 kDa cut-off filter to remove redox proteins such as ferredoxin, but no significant enhancement in Steady State Photocurrent or Spike Charge versus control measurements was observed with soluble fractions at $E_{app} \geq 0$ mV vs SHE (Figure S14c). Whilst this behavior could be caused by an electron mediator shuttling electrons between PSI and the electrode, cyclic voltammetry of soluble fraction revealed the presence of a small redox molecule with an $E_m$ of +53 mV vs SHE (Figure S15a), which is too positive to be responsible for the enhanced photocurrents observed at $E_{app}$ values < 0 mV vs SHE in the presence of soluble fractions. Fluorescence analysis revealed the presence of phycobilisomes in soluble fraction, alongside the presence of metabolites with blue fluorescence (Figure S15b). NADPH was shown to exhibit this same fluorescence peak (Figure S16a), with cyclic voltammetry showing that NADP(H) could not act as an electron mediator over the tested $E_m$ range (Figure S16b). Titration of NADPH into the electrolyte led to an enhancement of the Spike Charge at concentrations ≥ 500 µM (Figure S17a), suggesting the enhancements in photocurrents at lower $E_{app}$ values provided by soluble fractions may be caused by NADPH-dependent reduction of the P700 reaction center of PSI[26] (see Results for more information). The same effect was observed with $NADP^+$ addition (Figure S17b), consistent with the NADPH generation in the dark observed in spectroscopy experiments (Supporting Results 1, Figure S4f).

These data suggest that the Spike Charge of photocurrent profiles recorded under standard conditions (Fig. 1f) can be attributed to diffusible electron donors (as opposed to electron acceptors) being present in the sample, with the increase of the Spike Charge at high loadings (Figure S10a) being caused by the addition of electron donors trapped within the thylakoid membrane vesicles. This suggests electron transfer between the thylakoid membranes and the electrode is facilitated by DET.

Interestingly, the addition of soluble fraction and NADPH introduced a new feature in the photocurrent profile also- a transient decrease in the current occurring at light-dark transitions (Figure S20b). Further experiments are required to understand the molecular mechanisms underpinning this electrochemical signature.

#### Supporting Results 6: Cyclic voltammetry of thylakoid membranes

Cyclic voltammetry is an electrochemical technique in which the $E_{app}$ is scanned over time, giving rise to peaks in the current corresponding to the oxidation or reduction of different analytes. For any individual analyte which undergoes reversible electron transfer at the surface of the electrode, the midpoint between its oxidation peak ($E_{ox}$) and reduction peak ($E_{red}$) is equal to its $E_m$[41].

Cyclic voltammetry of thylakoid membranes performed in the absence and presence of DCMU and HQNO revealed a series of different oxidation and reduction peaks (Figure S18a), demonstrating that multiple thylakoid membrane redox cofactors were interfaced with the electrode. One pair of peaks, $E^1_{Ox}$ and $E^1_{Red}$ both exhibited a positive shift in potential in the presence of DCMU suggesting they originate from the same cofactor with an $E_m$ of +77 mV vs SHE (Figure S18b). $PQ(H_2)$ is known to have two distinct $E_m$ values depending on whether it is protein-bound within PSII as the cofactor $Q_B$ or free within the membrane PQ pool, with the latter having a more positive $E_m$[42]. Not only is the $E_m$ of this cofactor consistent with either $Q_B$ or $PQ(H_2)$[42], but the positive shift in potential in the presence of DCMU (and not HQNO) could be caused by displacement of $Q_B$ by the inhibitor, leading to its conversion into $PQ(H_2)$[43]. The presence of these redox peaks thereby demonstrates electron transfer between $PQ(H_2)$ within the thylakoid membrane and the electrode.

#### Supporting Results 7: Thylakoid membrane photocurrents from diverse species

Photocurrents were recorded using electrodes modified with thylakoid membranes from photosynthetic organisms other than *Synechocystis*. Thylakoid membranes of the cyanobacterium *Synechococcus elongatus* PCC 7942 produced identical photocurrent profiles to those of *Synechocystis*, albeit with differences in the magnitude of the Spike Charge and Steady State Photocurrent (Figure S21a). This demonstrates that thylakoid membrane electrochemistry is widely applicable for analysing cyanobacterial thylakoid membrane electron transport. Photocurrents were also recorded utilising thylakoids isolated from the model green algae *Chlamydomonas reinhardtii* and the plant *Nicotiana benthamiana*. Photocurrents could be reproducibly measured for both samples, which exhibited a monophasic photocurrent profile similar to those measured in other studies previously[44] (Figure S21b-c). The lack of a spike feature in these photocurrents could be due to the absence of RET in plant and algal thylakoid membranes[45]. However, further optimisation is required to ensure the lack of spike is not caused by inefficient wiring of thylakoid membranes to the electrode, which we suspect precluded measurements of Spike Charge in previous electrochemical studies of cyanobacterial thylakoid membranes[44]. Rapid photodegradation of these algal and plant thylakoid membranes was also observed in successive photocurrent measurements, suggesting further optimisation of the experimental conditions is required. Nonetheless, these results demonstrate that native membrane electrochemistry can be applied to analysing biological electron transport in membranes isolated from various organisms.

#### Supporting Discussion: Comparison with *in vivo* measurements of thylakoid membrane electron transport

Natural membrane electrochemistry relies on an *ex vivo* model system. To determine which phenomena this technique can investigate effectively, it is essential to compare the *ex vivo* and *in vivo* activities of the membranes being analyzed- cyanobacterial thylakoid membranes in this case.

Isolated thylakoid membranes were shown to retain their orientation and major membrane protein complexes, but much of the phycobilisome antennae detached from the surface of the membranes (Figure S2-3 and S15). Isolated thylakoid membranes retained activity of all LET chain components from PSII to PSI (Figure 2b-e and S4a-e), with PSII exhibiting acceptor-side limitation and PSI exhibiting donor-side limitation (Figure S4a-e)- as *in vivo*[46]. However, the isolation procedure led to damage of PSII (Figure S4a) as well as loss of soluble cytoplasmic proteins such as Ferredoxin and FNR (Figure S3 and S15). Ion transport across the membranes was similarly disrupted (Figure S4g and 10d), and non-photosynthetic NADPH synthesis was shown to occur (Figure S4f and S17). Functional electron transport between PSII and PSI was demonstrated (Figure 2g and S4c-d), although due to the non-specific or weak effects of available cytochrome $b_6f$ inhibitors[47,48], it proved difficult to implicate the role of this enzyme in electron transport between PSII and PSI. Plastoquinone was shown to be present in isolated thylakoid membranes (Figure S18), with the activity of respiratory dehydrogenases and oxidases being demonstrated in both electrochemical and spectroelectrochemical measurements (Figure 3c-f, S20 and S22). However, the presence of plasma membrane contamination in samples means some of these results may not be attributed to thylakoid membrane proteins. For example, measurements of terminal oxidase activity (Figure 3c-d) used a mutant lacking the ARTO protein, which is localized to the plasma membrane. Furthermore, whilst the presence of diffusible electron donors present in isolated thylakoid membrane samples was inferred (Figure S14-15), the identify of these metabolites was not determined. Consistency in photocurrent measurements was also observed across different abiotic conditions (Figure S10e) and when using thylakoid membranes isolated from different species (Figure S21).

It is important to note that the thylakoid membrane isolation used was selected for expediency and activity to facilitate the large number of electrochemical experiments performed herein. Lysis was performed by bead-vortexing, which was considered to be a gentler method than bead-beating or sonication. Thylakoid membrane purification methods used in proteomic experiments were tested[49], although these produced thylakoid membranes with no measurable photosynthetic activity. In recent years, new cyanobacterial thylakoid membrane isolation methods have been developed which may yield samples with higher purity and activity[50,51]. Future natural membrane electrochemistry experiments could use alternative membrane isolation methods which better retain the *in vivo* functionality of the membranes.

Nevertheless, the above assessment of our *ex vivo* model system is sufficient for investigating the electron transport activities of individual enzymes, or of electron transport chains given that they terminate at the cytoplasmic face of the membrane. Analysis of phycobilisome light harvesting or cytoplasmic proteins would require reconstitution of these proteins onto the surface of the membranes. Studies of proton motive force formation and dissipation, and its effect on electron transport, are probably unsuitable with this thylakoid isolation method due to the ion leak exhibited by the isolated membranes.

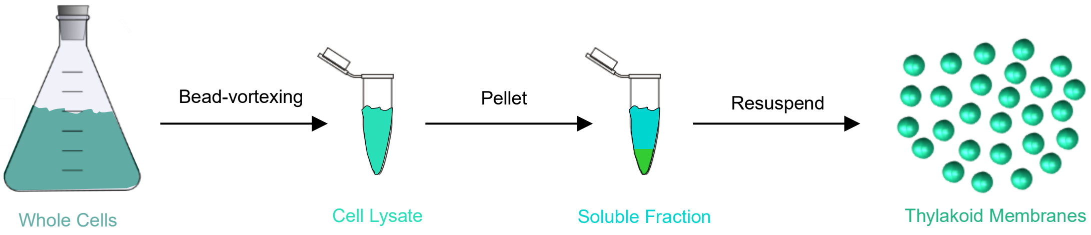


**Figure S1: Protocol for isolation of cyanobacterial thylakoid membrane vesicles.**

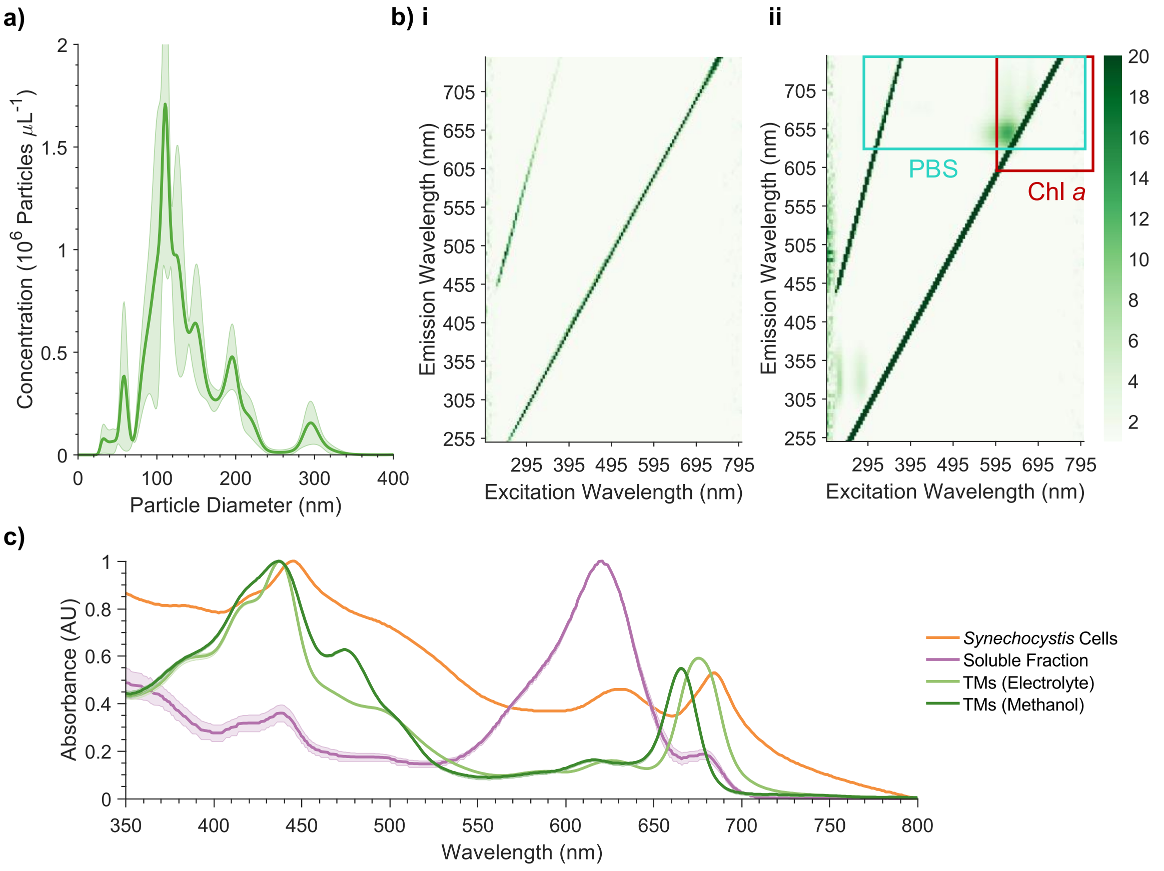


**Figure S2: Characterisation of *Synechocystis* thylakoid membrane vesicles.**
**a)** Size distribution of thylakoid membranes measured by NTA. **b)** Fluorescence excitation-emission matrix of **(i)** electrolyte buffer alone and **(ii)** supplemented with thylakoid membranes. **c)** Absorbance spectra of thylakoid membranes and other biological samples. NTA and absorbance data presented as the mean of biological replicates ± S.E.M. ($n$ = *3*). Other results ($n$ = 1). Chl *a*, chlorophyll *a*; PBS, phycobilisome; TM, thylakoid membrane.

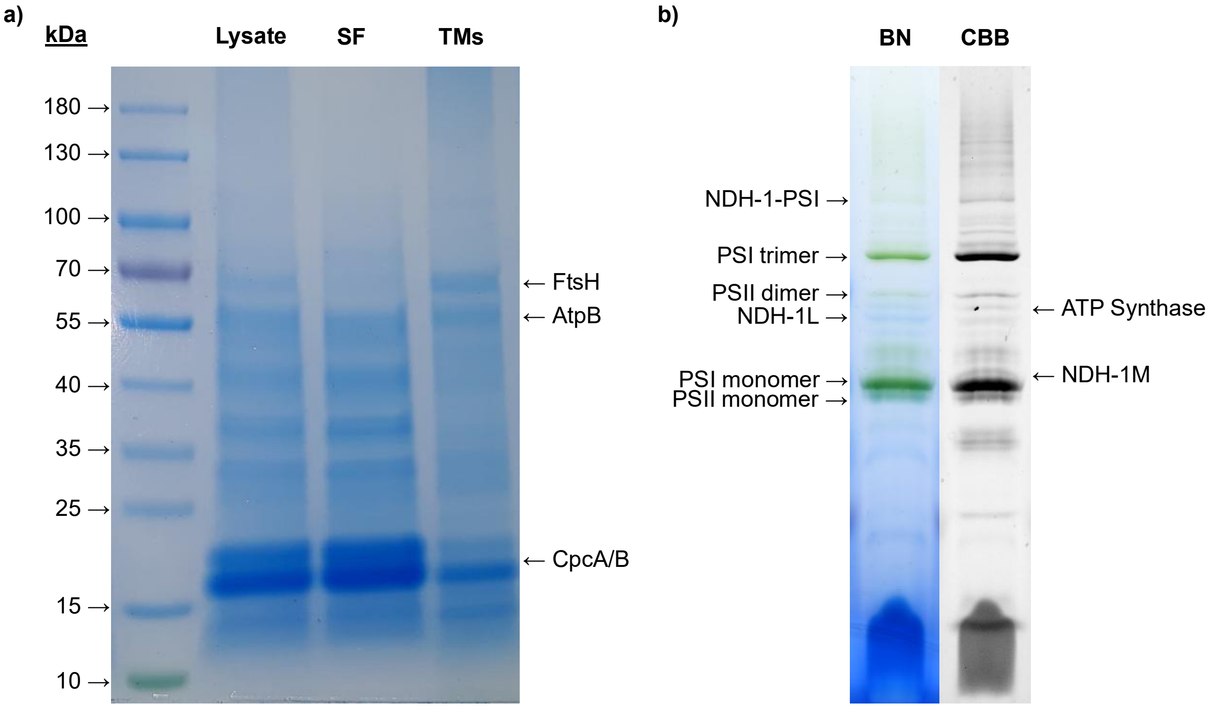


**Figure S3: PAGE analysis of isolated thylakoid membranes.**
**a)** SDS-PAGE and **b)** BN-PAGE of *Synechocystis* thylakoid membranes, with putative assignments of protein bands labelled[24,52]. ($n$ = 1). SF, soluble fraction; TM, thylakoid membrane.

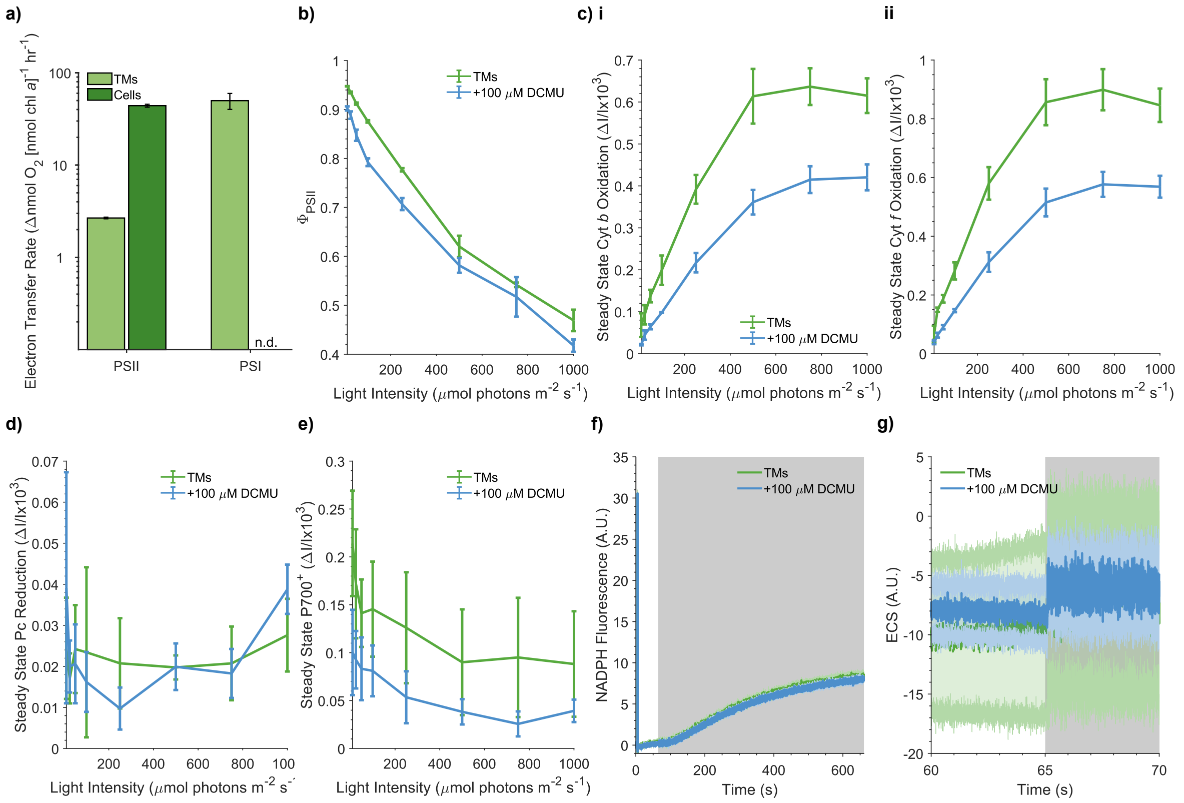


**Figure S4: Photosynthetic activity of isolated thylakoid membranes.**

**a)** Oxygen evolution rates performed under conditions to measure the maximal activity of PSII and PSI activity recorded under 1500 μmol $m^{-2}$ $s^{-1}$ of 680 nm light. PSII measurements were performed in the presence of 1 mM DCBQ and 1 mM ferricyanide as electron acceptors. PSI measurements were performed in the presence of 100 μM DCMU, 4 mM MV as an electron acceptor, and 20 mM ascorbate and 200 μM DCPIP as electron donors. **b)** $\phi_{PSII}$ and **c)** cytochrome *b*, and **d)** cytochrome *f* oxidation recorded at sequentially increasing intensities of 630 nm light, with 1 mM ferri-/ferrocyanide redox couple as an electron acceptor. **d)** Plastocyanin and **e)** P700 oxidation recorded at sequentially increasing intensities of 630 nm light, with 1 mM MV as an electron acceptor. **f)** NADPH fluorescence recorded at 500 μE with 1 mM $NADP^+$ as an electron acceptor. **g)** ECS measurements recorded at 500 μE with 1 mM ferri-/ferrocyanide redox couple as an electron acceptor. All spectroscopic measurements were performed with 630 nm light in the absence and presence of 100 μM DCMU, with chopped light (1 min on 1 min off), with the exception of NADPH fluorescence measurements (1 min on 10 min off). All data presented as the mean of biological replicates ± S.E.M. (*n* = *3*). Chl, chlorophyll; Cyt, cytochrome; DCMU, 3-(3,4-dichlorophenyl)-1,1-dimethylurea; ECS, electrochromic shift; NADP(H), nicotinamide adenine dinucleotide phosphate; Pc, plastocyanin; PSII, photosystem II; PSI, photosystem I; TM, thylakoid membrane.

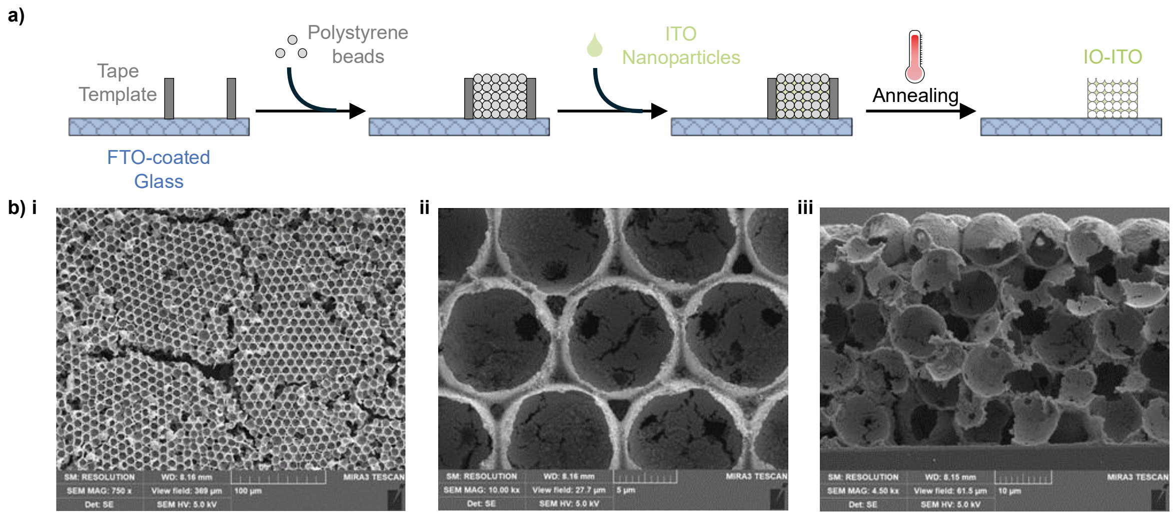


**Figure S5: Fabrication of IO-ITO electrodes.**
**a)** Protocol for IO-ITO electrode fabrication. **b)** SEM images of unmodified IO-ITO electrodes. **(i-ii)** Top views at two magnifications, and **(iii)** a cross section are shown. FTO, fluorine-doped tin oxide; IO-ITO, inverse opal-indium tin oxide.

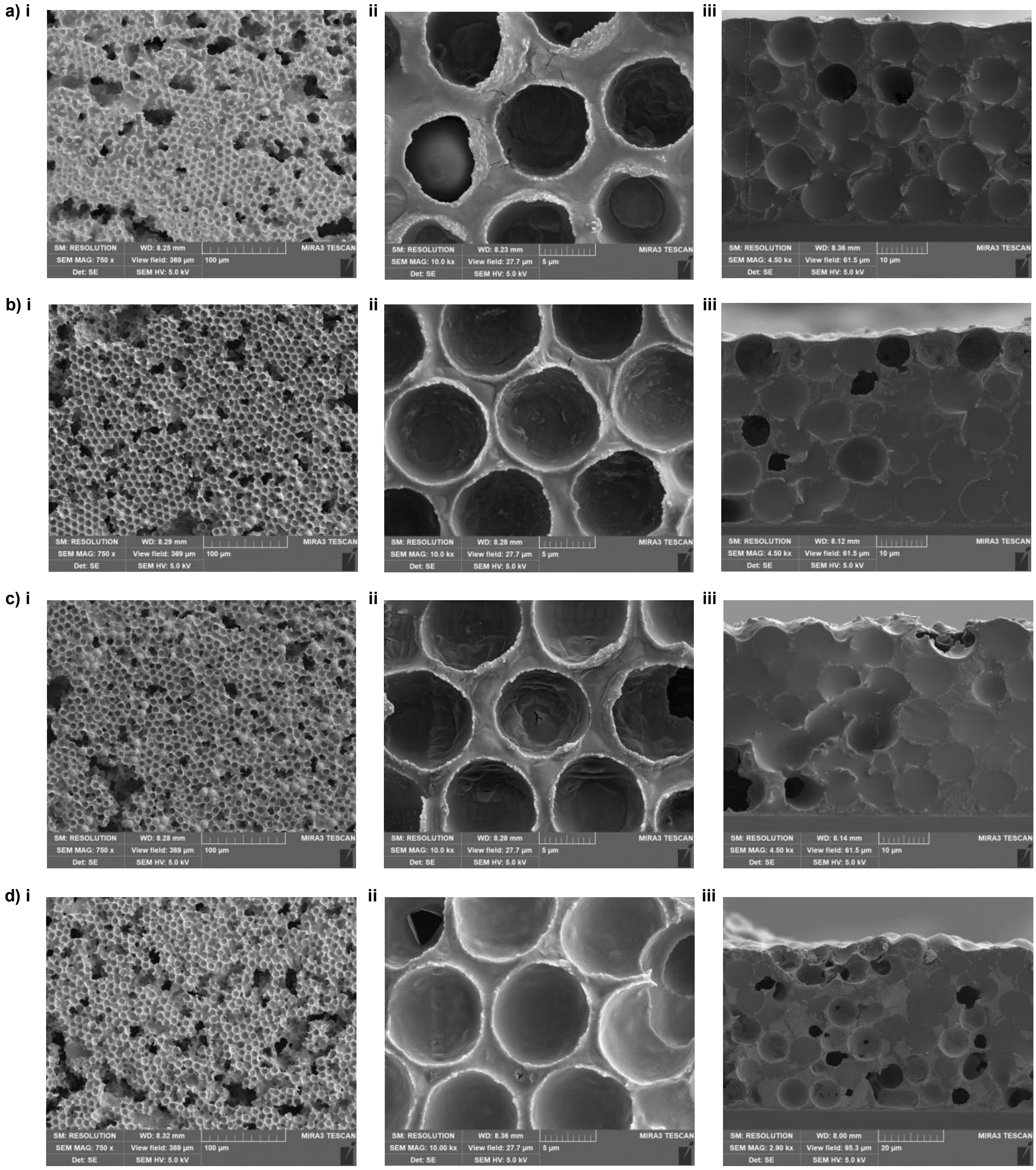


**Figure S6: Effect of thylakoid membrane loading on electrode structure.**
SEM images IO-ITO electrodes modified with thylakoid membranes with loadings of **a)** 2, **b)** 5, **c)** 10, and **d)** 20 μg chlorophyll *a*. **(i-ii)** Top views at two magnifications, and **(iii)** a cross section are shown. (*n = 1*). IO-ITO, inverse opal-indium tin oxide.

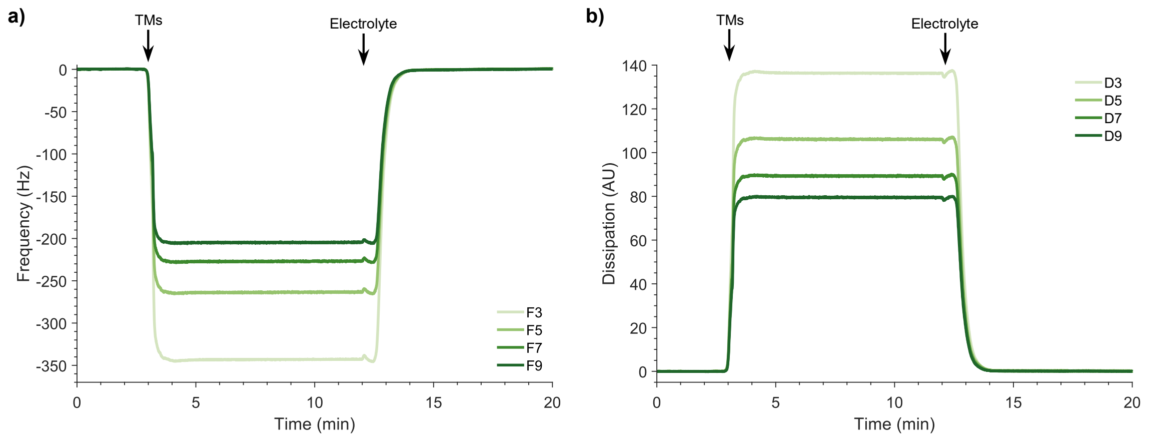


**Figure S7: QCM-D measurements of thylakoid membrane vesicles binding to ITO.** Simultaneous measurements of **a)** frequency and **b)** dissipation. Times at which thylakoid membranes and additional electrolyte buffer were added are labelled. (*n* = *1*). TM, thylakoid membrane.

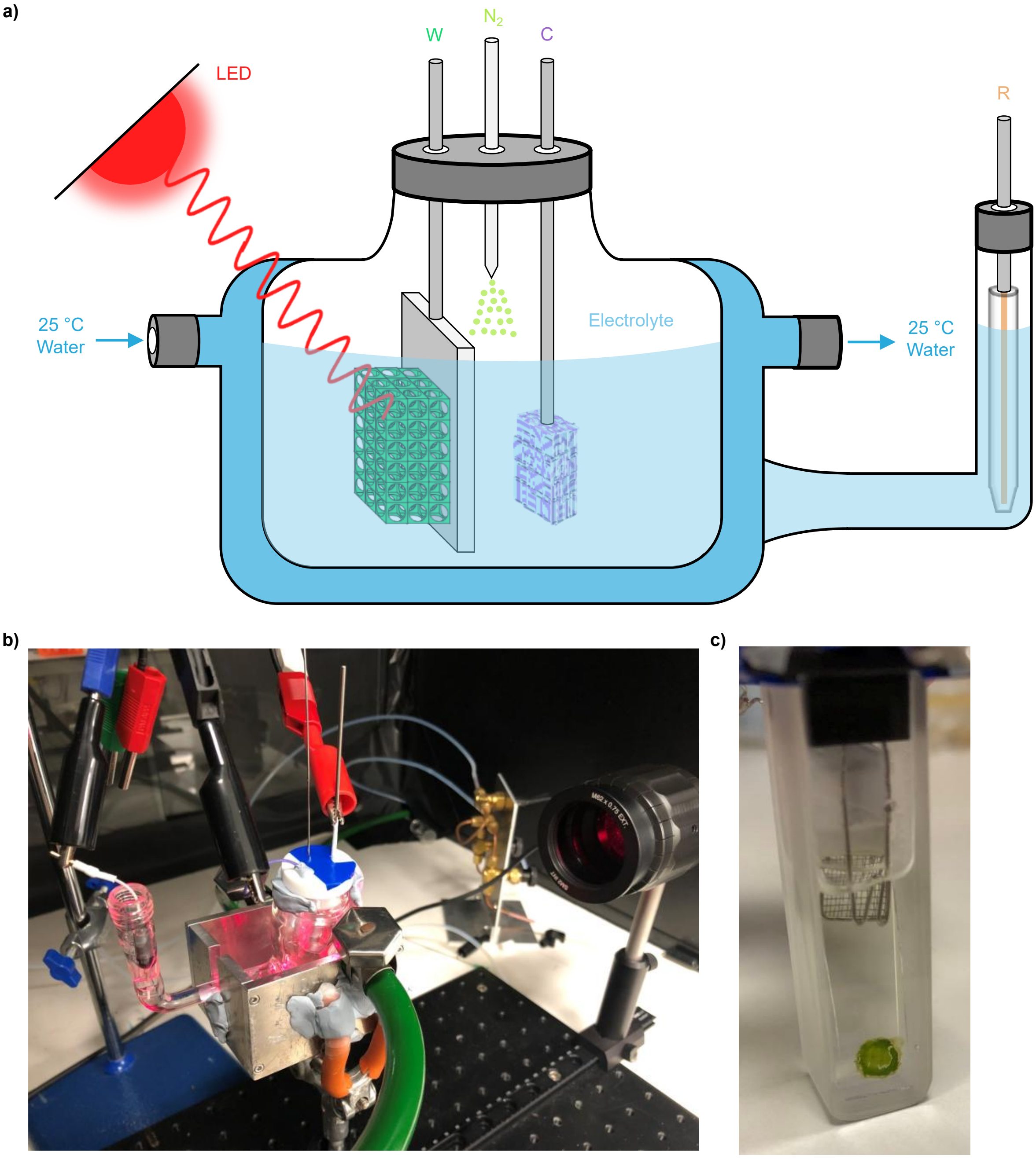


**Figure S8: Electrochemical apparatus for thylakoid membrane photoelectrochemistry.**
**a)** Schematic and **b)** photograph of the photo-bioelectrochemical cell used. **c)** Photograph of the spectroelectrochemical cell used. C, counter electrode; R, reference electrode; W, working electrode.

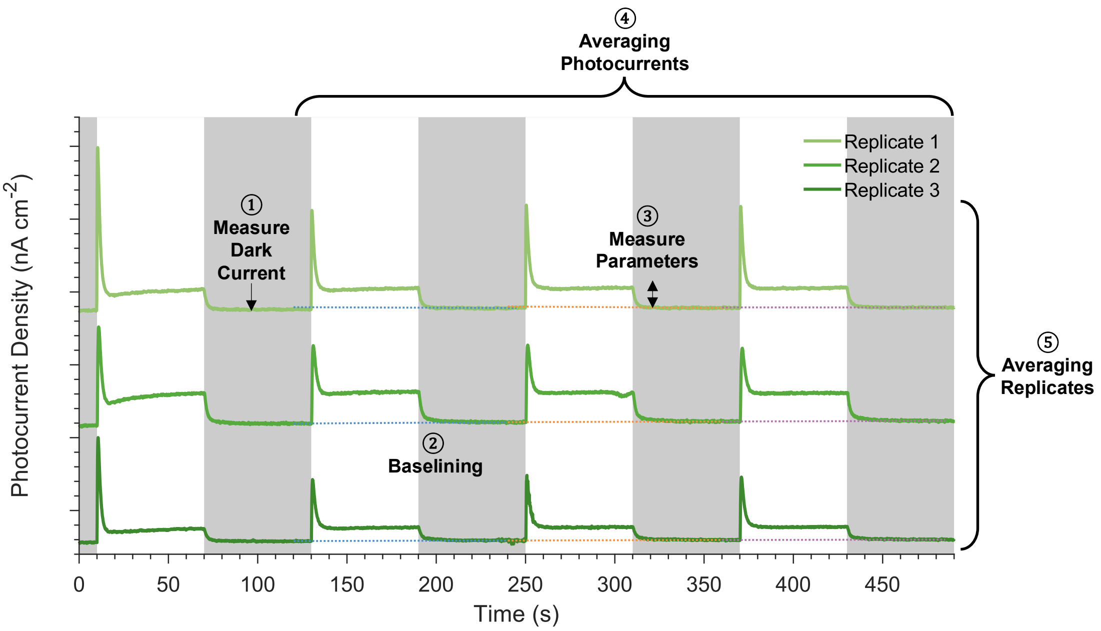


**Figure S9: Processing of thylakoid membrane chronoamperometry data.**
Chronoamperometry scans were recorded from multiple biological replicates under chopped light, with grey periods representing darkness and white periods light. Depicted scans were obtained from three biological replicates under standard conditions. Y-axis shown is a relative scale, with minor ticks representing 200 nA $cm^{-2}$. Data processing steps are labelled: ① dark current calculation, ② baselining, ③ parameter calculation, ④ averaging of successive photocurrent measurements, ⑤ averaging of biological replicates.

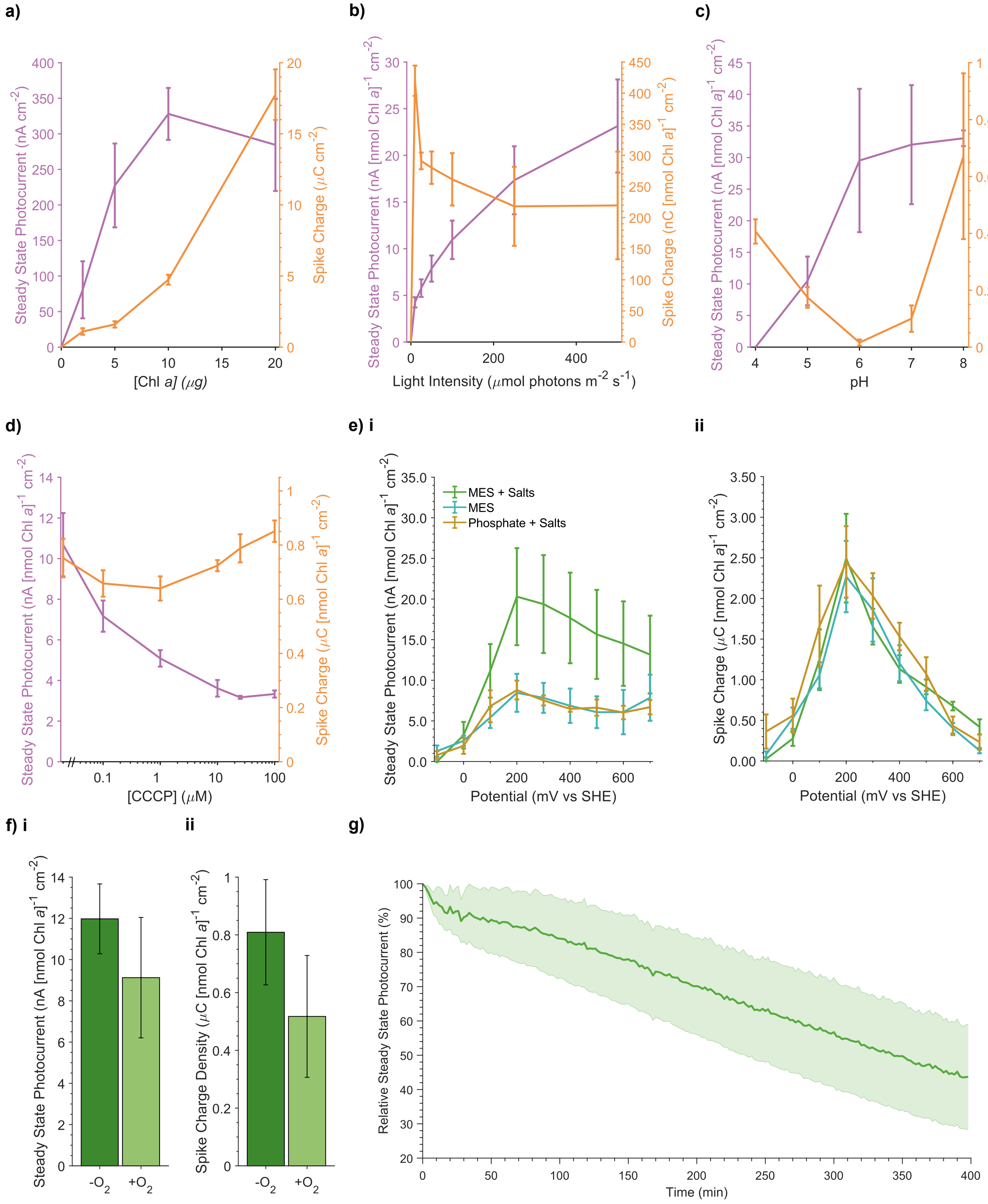


**Figure S10: Effect of experimental conditions on thylakoid membrane photocurrents.** Effect of **a)** chlorophyll a loading, **b)** light intensity, **c)** pH, and **d)** the ionophore CCCP on photocurrent parameters. **e)** Effect of electrolyte composition and $E_{app}$ on photocurrent parameters. Electrolyte was composed of either 50 mM MES or potassium phosphate, supplemented with or without salts (15 mM NaCl, 5 mM $MgCl_2$, 2 mM $CaCl_2$). **f)** Effect of the presence of oxygen on photocurrent parameters. Oxygen was removed through purging with $N_2$ gas. Sub-panels depict measurements of **(i)** Steady

State Photocurrent and **(ii)** Spike Charge magnitudes. **g)** Relative change in Steady State Photocurrent recorded from successive photocurrent measurements performed over 400 min. All data presented as the mean of biological replicates ± S.E.M. (*n* = *3*). *p*-values calculated using two-tailed unpaired *t*-tests (*** $p \le 0.001$, ** $0.001 < p \le 0.01$, $p \le 0.05$). Chl, chlorophyll; CCCP, Carbonyl cyanide m-chlorophenyl hydrazone; DCMU, 3-(3,4-dichlorophenyl)-1,1-dimethylurea; MES, 2-(N-morpholino)ethanesulfonic acid; SHE, standard hydrogen electrode.

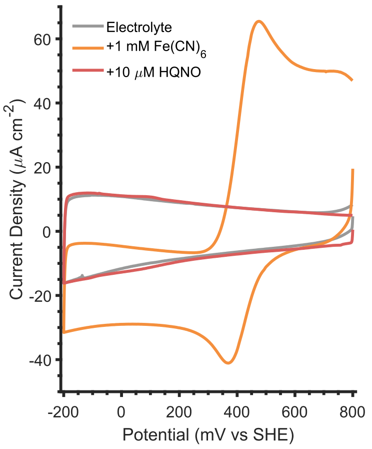


**Figure S11: Cyclic voltammetry of redox active compounds.**
Cyclic voltammograms of 1 mM ferri-/ferrocyanide and 10 μM HQNO in electrolyte buffer. Experiments were performed with an IO-ITO working electrode, platinum mesh counter electrode and Ag/AgCl reference electrode, using a scan rate of 2 mV $s^{-1}$. (*n = 1*). $Fe(CN)_6^{3-}/Fe(CN)_6^{4-}$, ferri-/ferrocyanide redox couple; HQNO, N-oxo-2-heptyl-4-Hydroxyquinoline; SHE, standard hydrogen electrode.

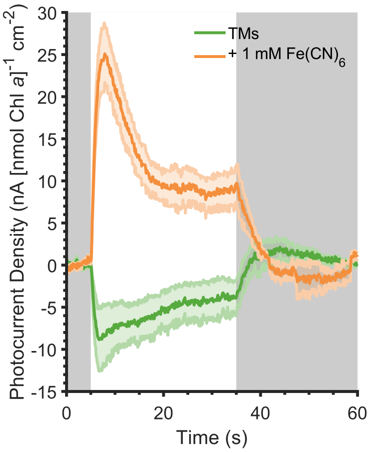


**Figure S12: Effect of ferrocyanide on thylakoid membrane photocurrents.**
Baselined thylakoid membrane photocurrent profiles recorded at an $E_{app}$ of -600 mV vs SHE. Data presented as the mean of biological replicates ± S.E.M. (*n* = *3*). Data same as in Figure 2c. $Fe(CN)_6$, ferrocyanide.

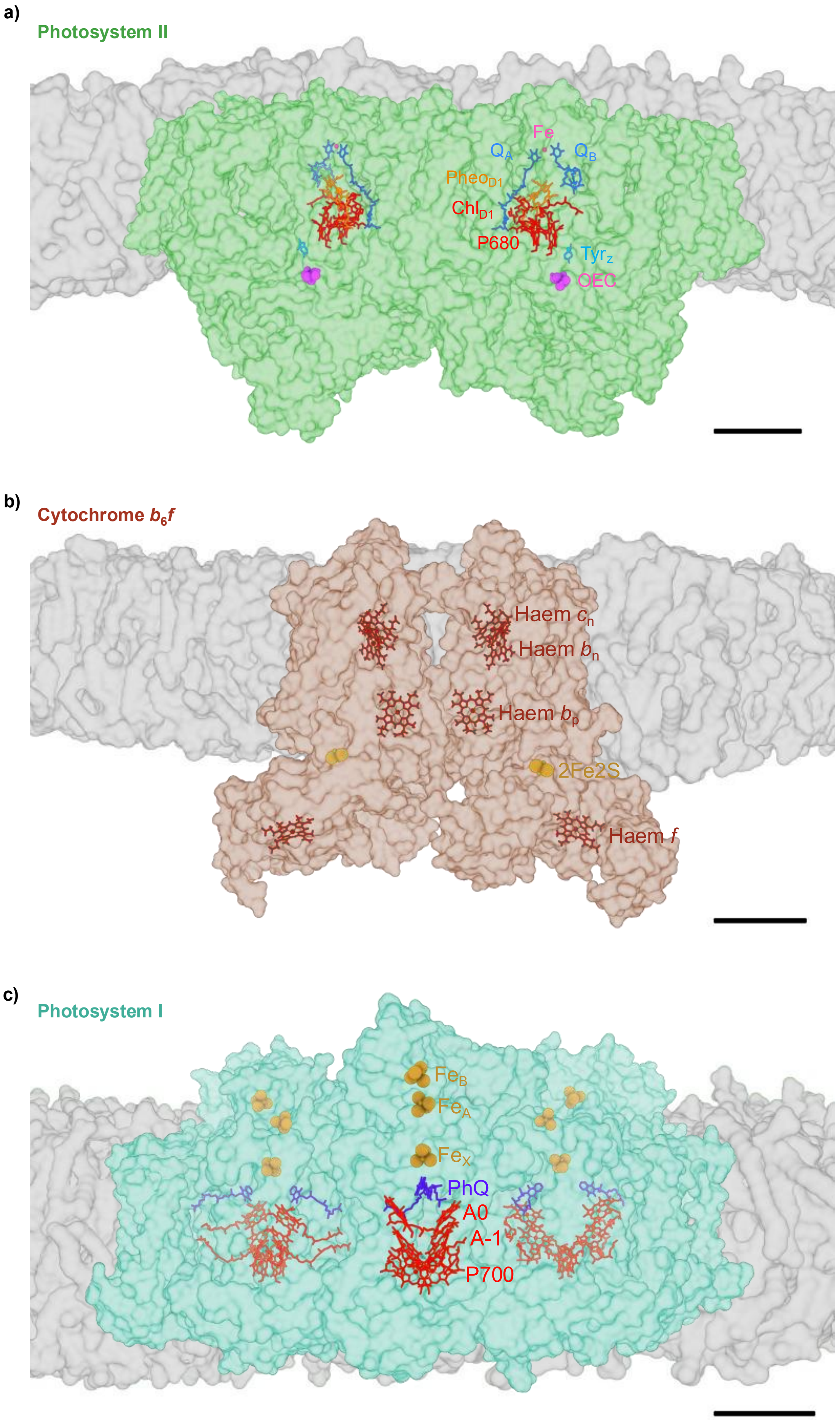
a)
Photosystem II
Fe
$Q_A$
$Q_B$
$Pheo_{D1}$
$Chl_{D1}$
P680
$Tyr_Z$
OEC
b)
Cytochrome $b_6f$
Haem $c_n$
Haem $b_n$
Haem $b_p$
2Fe2S
Haem $f$
c)
Photosystem I
$Fe_B$
$Fe_A$
$Fe_X$
PhQ
A0
A-1
P700

**Figure S13: Structural analysis of thylakoid membrane protein complexes embedded in a lipid bilayer.**

Structural models of **a)** photosystem II dimer (PDB: 3WU2[20]), **b)** cytochrome $b_6f$ dimer (PDB: 4H44[21]), and **c)** photosystem I trimer (PDB: 1JBO[22]). Insertion of these biological assemblies into lipid bilayers was predicted by molecular dynamics simulations, as part of the MemProtDB resource[23]. Front views are shown. Scale bar: 2nm. A, acceptor; Chl, chlorophyll; Fe, iron-sulphur cluster; 2Fe2S, Rieske iron-sulphur cluster; OEC, oxygen evolving complex; P, primary donor; Ph, phylloquinone; Pheo, pheophytin; Q, quinone; Tyr, tyrosine.

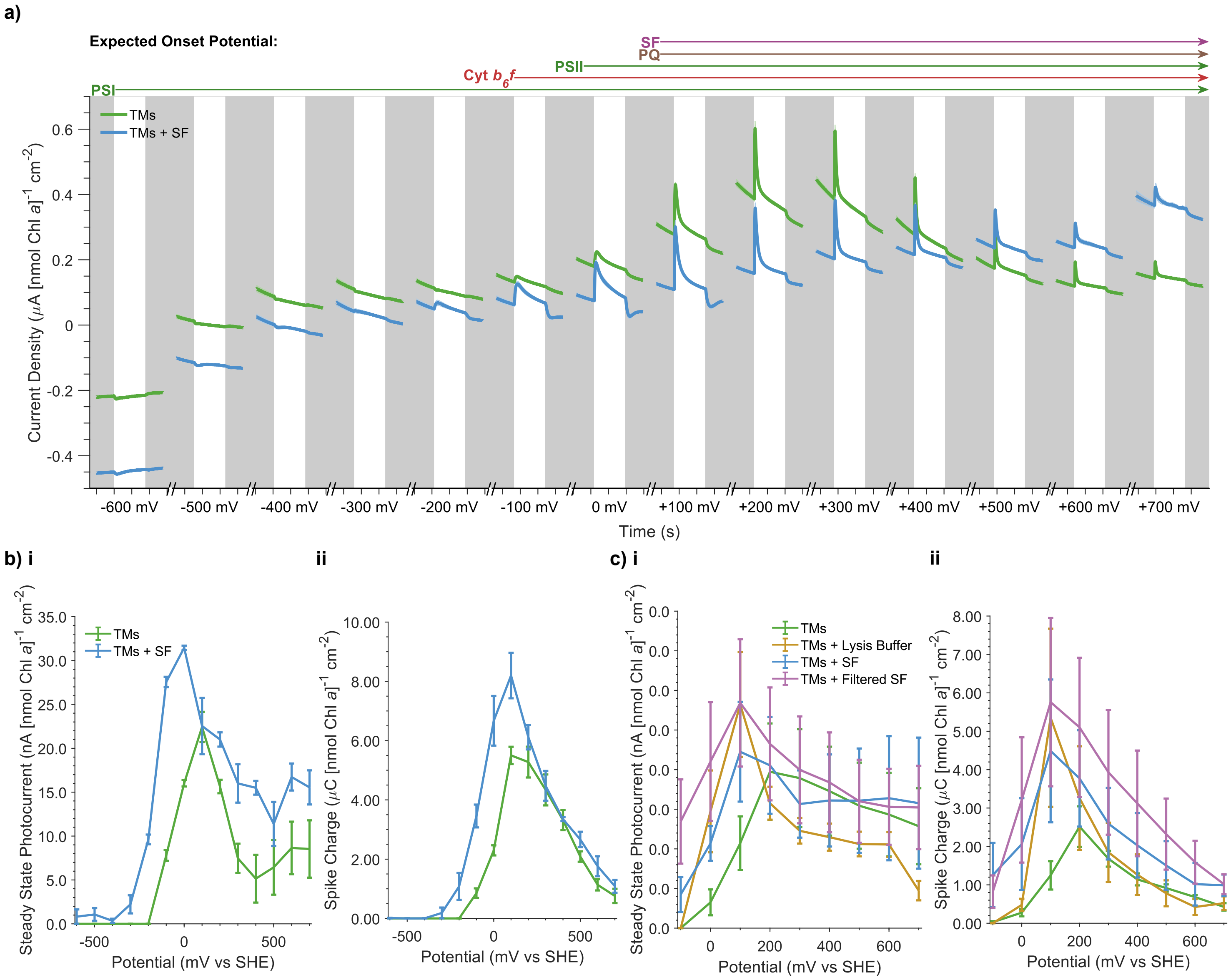


**Figure S14: Effect of soluble proteins and metabolites on thylakoid membrane photocurrents.** **a)** Stepped chronoamperometry scan of thylakoid membranes, with photocurrents recorded at increasing $E_{app}$ values (mV vs SHE) and in the absence and presence of 1 mM $Fe(CN)_6$ along with an enzymatic oxygen removal systems (1 mM glucose, 100 μg $mL^{-1}$ glucose oxidase, 50 μg $mL^{-1}$ catalase). **b)** Photocurrent parameters, calculated from **a.** **c)** Effect of lysis buffer, soluble fraction and 3 kDa-filtered soluble fraction on photocurrent parameters, recorded at increasing $E_{app}$ values in the absence of enzymatic oxygen removal. Sub-panels depict measurements of **(i)** Steady State Photocurrent and **(ii)** Spike Charge magnitudes. Data presented as the mean of biological replicates ± S.E.M. (*n* = *3*). Chl, chlorophyll; Cyt $b_6f$, cytochrome $b_6f$; FSF, 3 kDa-filtered soluble fraction; PQ, plastoquinone; PSII, photosystem II; PSI, photosystem I; SF, soluble fraction; SHE, standard hydrogen electrode; TM, thylakoid membrane.

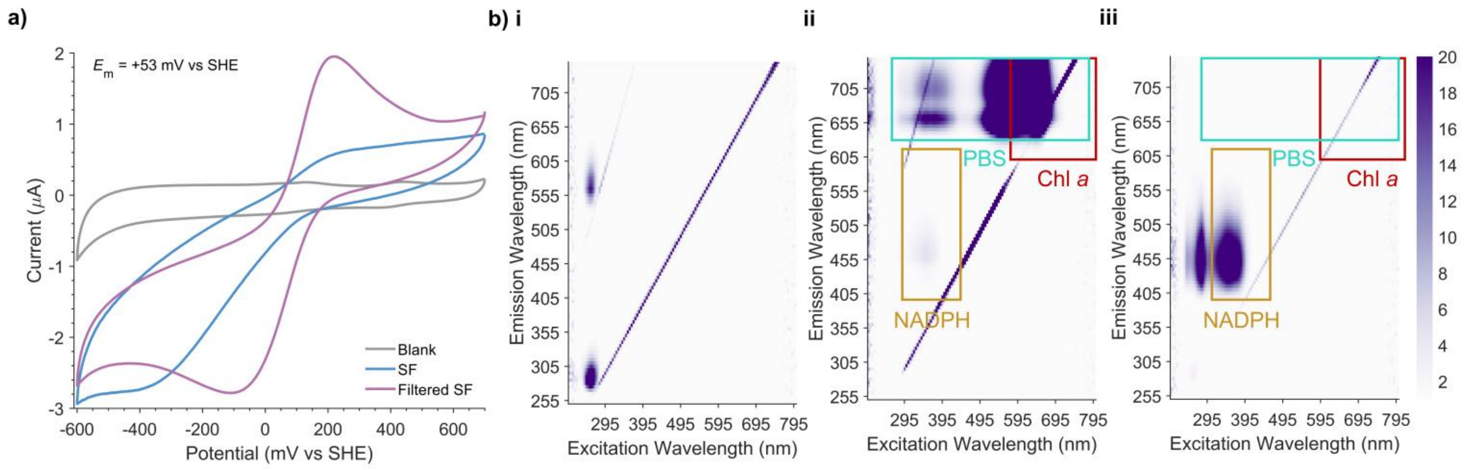


**Figure S15: Composition of soluble fractions.**
**a)** Cyclic voltammetry of soluble fraction and 3 kDa-filtered soluble performed with an ITO-coated glass working electrode, platinum mesh counter electrode and Ag/AgCl reference electrode with a scan rate of 10 mV $s^{-1}$. **d)** Fluorescence excitation emission matrices of **(i)** lysis buffer, **(ii)** soluble fraction, and **(iii)** 3 kDa-filtered soluble fraction. (*n* = *1*). Chl, chlorophyll; NADP(H), nicotinamide adenine dinucleotide phosphate; PBS, phycobilisomes; SF, soluble fraction; SHE, standard hydrogen electrode.

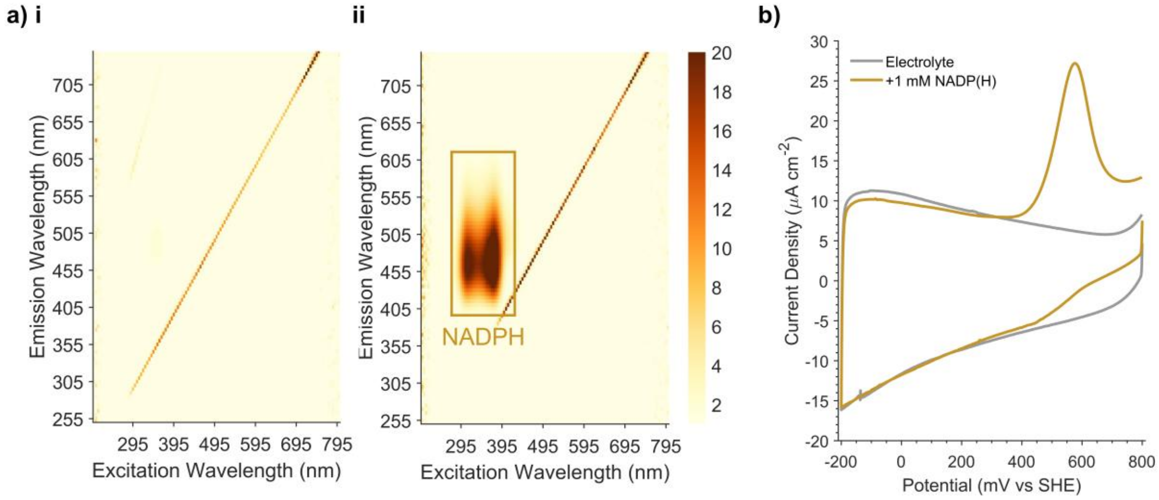


**Figure S16: Characterisation of NADP(H) solutions.**
**a)** Fluorescence excitation emission matrices of 5 mM **(i)** $NADP^+$ and **(ii)** NADPH in electrolyte buffer. (*n = 1*). **b)** Cyclic voltammogram of NADP(H), performed with an IO-ITO working electrode, platinum mesh counter electrode and Ag/AgCl reference electrode, in electrolyte buffer supplemented with 2.5 mM of $NADP^+$ and 2.5 mM of NADPH, using a scan rate of 2 mV $s^{-1}$. (*n = 1*). NADP(H), nicotinamide adenine dinucleotide phosphate; SHE, standard hydrogen electrode.

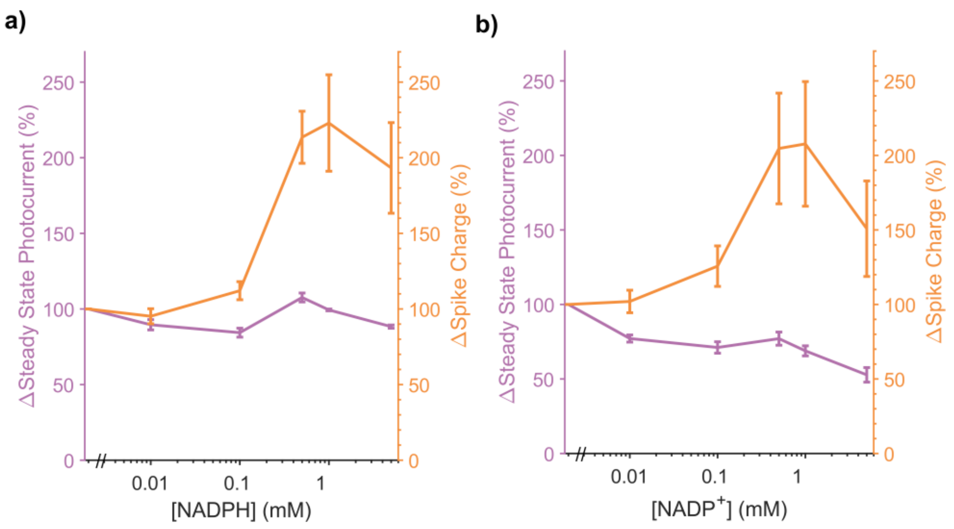


**Figure S17: Effect of NADP(H) on thylakoid membrane photocurrents.**
Effect of **a)** NADPH and **b)** $NADP^+$ addition on relative photocurrent parameters. Sub-panels depict measurements of **(i)** Steady State Photocurrent and **(ii)** Spike Charge magnitudes. Data presented as the mean of biological replicates ± S.E.M. (*n = 3*). NADP(H), nicotinamide adenine dinucleotide phosphate; SHE, standard hydrogen electrode.

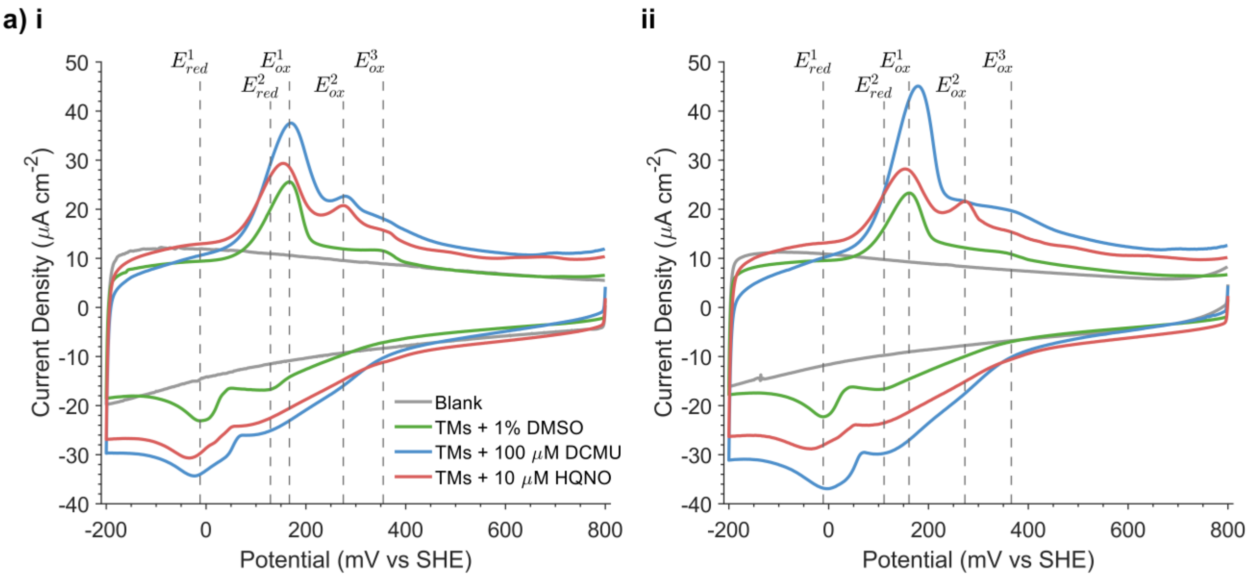


**b)**

| Peaks (mV vs SHE) | +DMSO D | +DMSO L | +DCMU D | +DCMU L | +HQNO D | +HQNO L |
|---|---|---|---|---|---|---|
| $^{\dagger}E^1_{red}$ | -12 | -11 | -24 | -2 | -34 | -36 |
| $^{\ddagger}E^2_{red}$ | +129 | +111 | +129 | +117 | +129 | +111 |
| $^{\dagger}E^1_{ox}$ | +167 | +161 | +170 | +179 | +167 | +153 |
| $^{\dagger}E^2_{ox}$ | / | / | +279 | / | +275 | +273 |
| $^{\ddagger}E^3_{ox}$ | +355 | / | / | / | +366 | +367 |

$Q_B$ / $PQ(H_2)$ [$E_m$ = ~+77 mV]

**Figure S18: Identifying redox cofactors with cyclic voltammetry.**
**a)** Cyclic voltammograms of thylakoid membranes recorded in electrolyte buffer supplemented with DMSO (1 %), DCMU (100 μM) and HQNO (10 μM), alongside blank measurements recorded without thylakoid membranes. Experiments were performed with an IO-ITO working electrode, platinum mesh counter electrode and Ag/AgCl reference electrode, using a scan rate of 2 mV $s^{-1}$. Identifiable oxidation and reduction peaks are labelled. Sub-panels depict measurements recorded **(i)** in darkness and **(ii)** under 50 μmol $m^{-2}$ $s^{-1}$ of 680 nm light. **b)** The potentials of oxidation and reduction peaks measured from **a**. Potentials measured through a peak search algorithm are denoted with †, whilst potentials measured manually are denoted with ‡. Peaks $E^1_{Red}$ and $E^1_{Ox}$ are consistent with a redox couple for $Q_B$ in PSII and/or $PQ(H_2)$[42]. D, dark; DCMU, 3-(3,4-dichlorophenyl)-1,1-dimethylurea; DMSO, dimethyl sulfoxide; HQNO, N-oxo-2-heptyl-4-Hydroxyquinoline; L, light; SHE, standard hydrogen electrode; TM, thylakoid membrane.

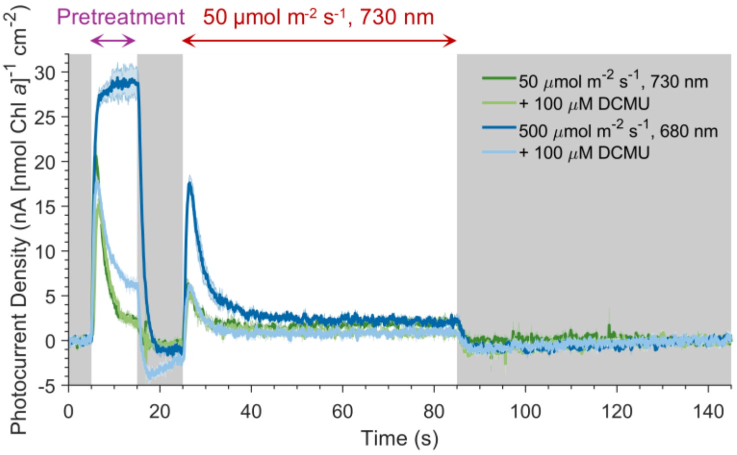


**Figure S19: Effect of light pre-treatment conditions on thylakoid membrane photocurrents.** Thylakoid membrane photocurrent profiles recorded following different light pre-treatments. Data presented as the mean of biological replicates ± S.E.M. (*n* = *3*). Photocurrent parameters in Figure 2g. Chl, chlorophyll; DCMU, 3-(3,4-dichlorophenyl)-1,1-dimethylurea.

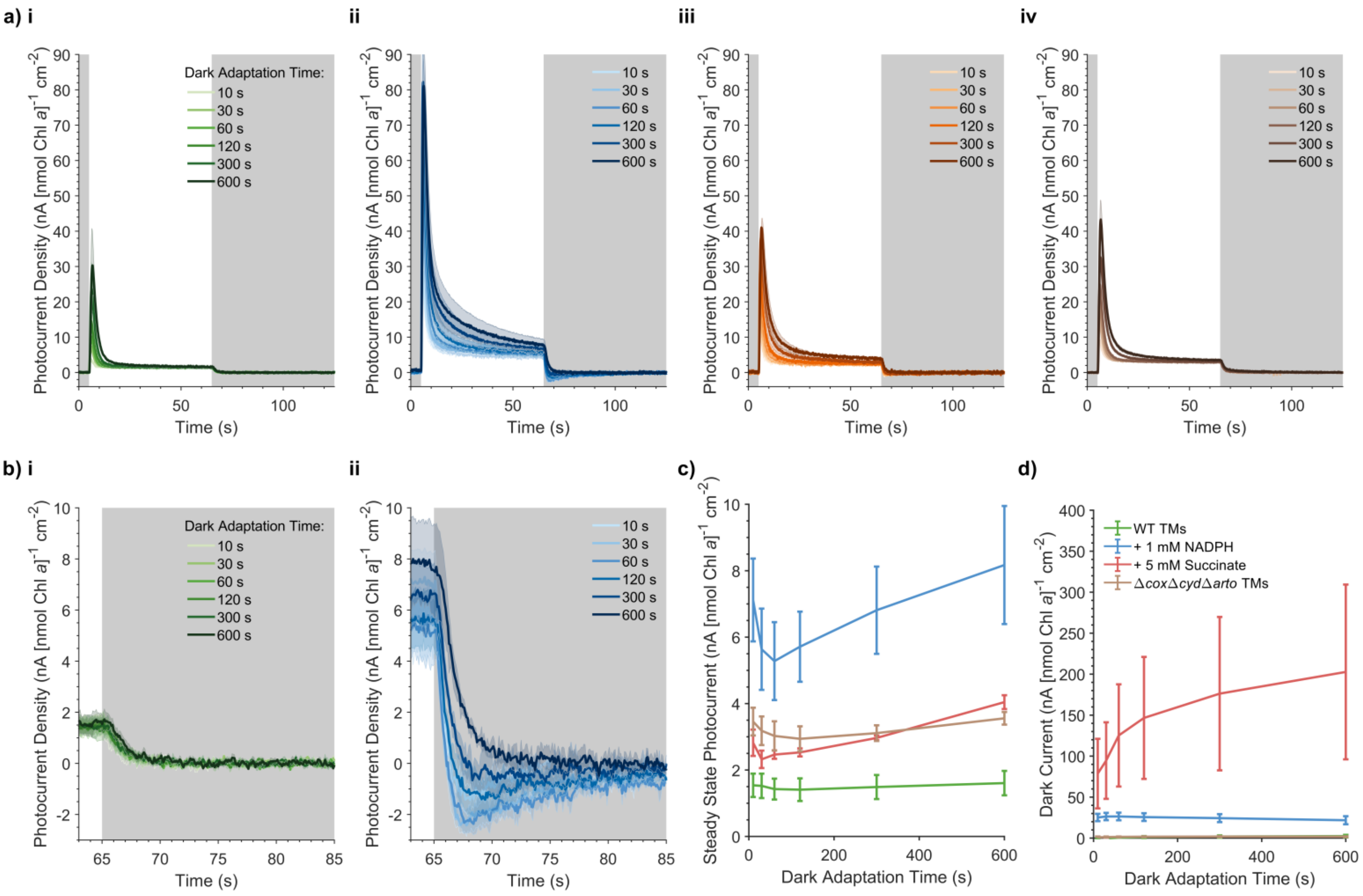


**Figure S20: Effect of plastoquinone pool reduction on thylakoid membrane photocurrents.** **a)** Thylakoid membrane photocurrent profiles recorded in increasing dark adaptation times and the **(i)** absence (same data as Figure 3ci) and presence of **(ii)** 1 mM NADPH, **(iii)** 5 mM Succinate, or **(iv)** using thylakoid membranes isolated from from Δ*cyd*Δ*cox*Δ*arto* cells. **b)** Same data as panel ai-ii, showing a zoom in on the dark-light transition. **c)** Steady State Photocurrents and **d)** dark currents recorded at all dark adaptation times. Data presented as the mean of biological replicates ± S.E.M. (*n* = *3*). Companion to Figure 3c. Chl, chlorophyll; NAD(P)(H), nicotinamide adenine dinucleotide (phosphate); TM, thylakoid membrane; WT, wild type.

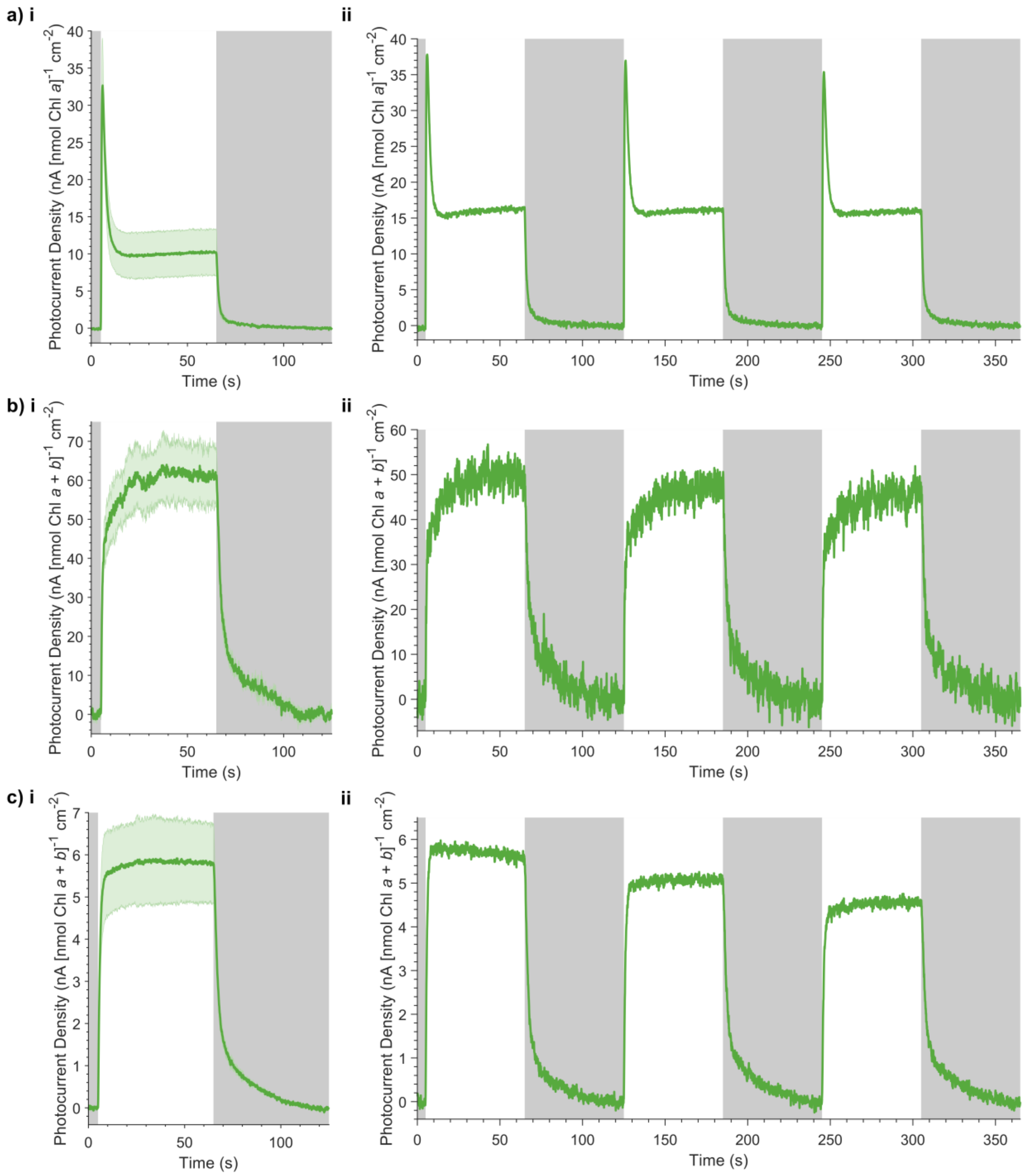


**Figure S21: Photocurrent measurements of thylakoid membranes from different species.** Photocurrents of thylakoid membranes isolated from **a)** *S. elongatus*, **b)** *C. reinhardtii* and **c)** *N. benthamiana*. Sup-panels depict **(i)** the mean of biological replicates ± S.E.M. (*n* = *3*), and **(ii)** representative chronoamperometry scans, depicting successive photocurrent measurements. (*n* = *1*). Chl, chlorophyll.

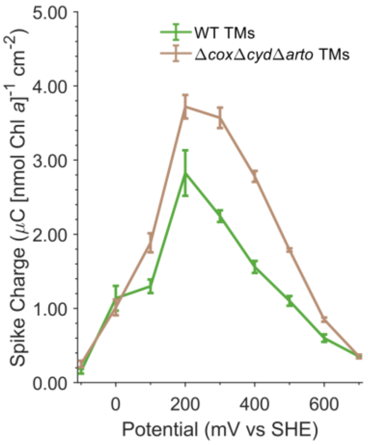


**Figure S22: Effect of terminal oxidases on thylakoid membrane photocurrents.**
Absolute measurements of Spike Charge recorded from photocurrents obtained at $E_{app}$ values ≥ +200 mV vs SHE for thylakoid membrane from wild type and Δ*cyd*Δ*cox*Δ*arto* without enzymatic oxygen removal. Same data as in Figure 3d. Chl, chlorophyll; SHE, standard hydrogen electrode; TM, thylakoid membrane; WT, wild type.

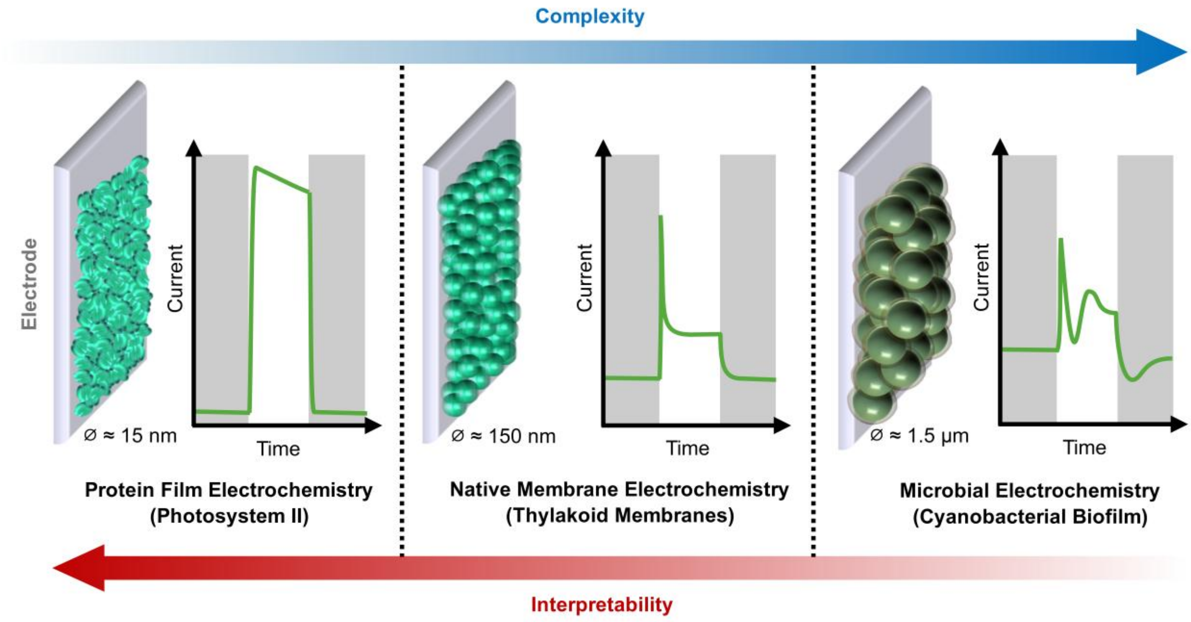


**Figure S23: Comparison of bioelectrochemical methods.**
Schematic showing electrochemical measurements obtained by protein electrochemistry (Photosystem II), native membrane electrochemistry (thylakoid membranes), and microbial electrochemistry (cyanobacterial biofilms). An approximate diameter is shown for each analyte. Representations of the photocurrent profile are depicted, showing trade-offs between the complexity and interpretability of these data[53].

| Pathway | Figures |
|---|---|
| ① | 2b-f, S10b-d, S13a |
| ② | S20d |
| ③ | 3d, S18 |
| ④ | 3d, S13b |
| ⑤ | 2b, 2f |
| ⑥ | 2b-f, 3b-f, S10b-d, S13c |

**Table S1: Evidence for thylakoid membrane and interfacial electron transfer pathways.** Companion to Figure 4.
